# MCM complexes are barriers that restrict cohesin-mediated loop extrusion

**DOI:** 10.1101/2020.10.15.340356

**Authors:** Bart J. H. Dequeker, Hugo B. Brandão, Matthias J. Scherr, Johanna Gassler, Sean Powell, Imre Gaspar, Ilya M. Flyamer, Wen Tang, Roman Stocsits, Iain F. Davidson, Jan-Michael Peters, Karl E. Duderstadt, Leonid A. Mirny, Kikuё Tachibana

**Author notes:** These authors contributed equally to this work (Hugo B. Brandão, Matthias J. Scherr).

## Abstract

Eukaryotic genomes are compacted into loops and topologically associating domains (TADs), which contribute to transcription, recombination and genomic stability. Cohesin extrudes DNA into loops that are thought to lengthen until CTCF boundaries are encountered. Little is known about whether loop extrusion is impeded by DNA-bound macromolecular machines. We demonstrate that the replicative helicase MCM is a barrier that restricts loop extrusion in G1 phase. Single-nucleus Hi-C of one-cell embryos revealed that MCM loading reduces CTCF-anchored loops and decreases TAD boundary insulation, suggesting loop extrusion is impeded before reaching CTCF. Single-molecule imaging shows that MCMs are physical barriers that frequently constrain cohesin translocation *in vitro.* Simulations are consistent with MCMs as abundant, random barriers. We conclude that distinct loop extrusion barriers contribute to shaping 3D genomes.

**One Sentence Summary:** MCM complexes are obstacles that impede the formation of CTCF-anchored loops.

---

The genome of eukaryotes is folded into loops that are generated by structural maintenance of chromosomes (SMC) proteins [reviewed in (1)]. Structures emerging as a result of loop extrusion are detected by chromosome conformation capture (Hi-C) experiments, and are generated by a mechanism that requires the cohesin holocomplex, consisting of Smc1-Smc3-Scc1 with Nipbl and Stag1/Stag2 (2–4). Single cohesin complexes, similarly to condensin complexes, can actively form DNA loops *in vitro* (5–8). The extrusion process is hypothesized to form progressively larger loops until cohesin encounters a barrier and/or is released by Wapl (9–13). The predominant barrier to loop extrusion in vertebrates is the CTCF zinc finger transcription factor. CTCF bound to motifs in convergent orientation is required for formation of CTCF-anchored loops and generates insulation at the boundaries of TADs (9,14). Extrusion barriers, such as CTCF, play an instructive role in establishing extrusion-mediated structures visible in Hi-C, as evident from CTCF-depletion experiments that results in disappearance of major hallmarks of Hi-C maps (4,15). However, the loop extrusion machinery likely encounters other obstacles on chromatin, such as nucleosomes, RNA polymerases and other macromolecular complexes and enzymes. While RNA polymerases were shown to be moving barriers for condensin translocation in bacteria (16) and affecting cohesin translocation in eukaryotes (17,18), it remains unknown how SMCs can extrude loops on busy eukaryotic DNA bound by myriads of other proteins. Moreover, whether and how other DNA-bound macromolecular machines can influence 3D genome architecture in eukaryotes is not known and can be critical for understanding their function.

The minichromosome maintenance (MCM) complex is an abundant macromolecular machine that is essential for DNA replication in eukaryotes and archaea (19). MCM2-7 complexes are loaded at replication origins by the origin recognition complex (ORC), Cdc6 and Cdt1 to form the prereplication complex (pre-RC) during late mitosis and G1 phase (20). The head-to-head double MCM hexamer topologically entraps double-stranded DNA and is catalytically inactive as a helicase until initiation of DNA replication (21). Interestingly, 10-100-fold more MCMs are loaded onto chromatin than are needed for normal S phase progression. This is referred to as the “MCM paradox” (22). One hypothesis to explain this phenomenon is that the surplus complexes mark dormant origins that fire under conditions such as DNA damage checkpoint activation (23). Whether they have any functional consequences in unperturbed S or even G1 phase remains unclear. Given MCM abundance, the long residence time on chromatin (>6 h) (24) and size (13 nm) (25) comparable to FtsK helicase (12.5 nm) (fig. S1) that can push cohesin on DNA *in vitro* (26), we asked whether MCMs can act as obstacles to cohesin-dependent loop extrusion and in this way influence genome architecture.

To test this hypothesis, we utilized the oocyte-to-zygote transition (OZT) to investigate whether MCM loss affects cohesin-mediated loop extrusion, CTCF-anchored loops, and TADs. Oocytes are female germ cells that undergo the meiosis I division, complete the meiosis II division upon fertilization and generate one-cell embryos (zygotes). These harbour spatially separate maternal and paternal pronuclei whose chromatin is organized into cohesin-dependent loops and TADs (2,28). While cells of the OZT are limited by paucity of material, they offer key advantages including the ability: 1) to study MCM loading on *de novo* assembled paternal chromatin occurring after fertilization in G1 phase, 2) to decipher effects on chromatin organization in a haplotype-resolved manner, 3) to manipulate pre-RC assembly without interfering with meiotic cell cycle progression, because there is no DNA replication between meiosis I and II, and 4) to disentangle direct from indirect effects via gene expression changes, since mature oocytes and G1 zygotes are transcriptionally inactive (27).

To generate zygotes deficient in MCMs, we interfered with the Cdt1-mediated loading pathway. Cdt1 deposits MCMs onto chromatin, a reaction that is inhibited by geminin, a target of the anaphase-promoting complex/cyclosome (APC/C) (fig. S2A) (29). Mutation of geminin’s destruction box generates a non-degradable version (geminin^L26A^) that inhibits Cdt1-mediated MCM loading in G1 phase (fig. S2A) (30). To achieve this, mature germinal vesicle (GV)-stage mouse oocytes were microinjected with mRNA encoding an injection marker GFP with or without geminin^L26A^ (Fig. 1A). Oocytes resumed meiosis I, progressed to metaphase II and were fertilized *in vitro* (IVF) to generate zygotes. Following extraction of soluble proteins, fixation and immunofluorescent staining, little or no chromatin-associated MCMs were detected in G1 phase zygotes expressing geminin^L26A^ (referred to as ‘MCM loss’) (Fig. 1B and fig. S2, B and D), demonstrating efficient inhibition of Cdt1-mediated MCM loading.

**Fig. 1.**
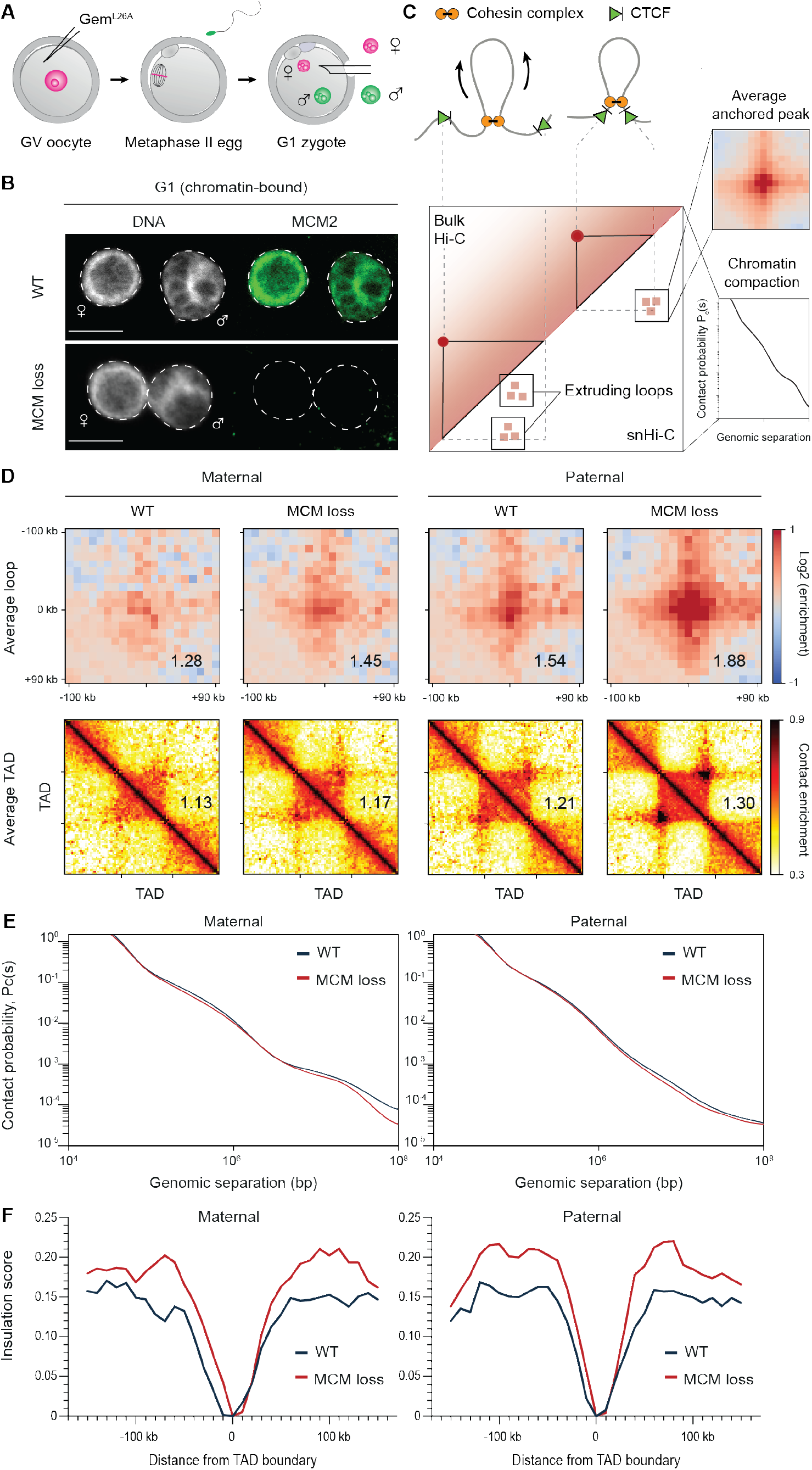
Chromatin-bound MCM impedes loop and TAD formation in G1 phase zygotes. **(A)** To prevent MCM recruitment to DNA, GV oocytes were injected with geminin^L26A^ mRNA, *in vitro* matured to meiosis II, followed by *in vitro* fertilization to form zygotes. Maternal (magenta) and paternal (green) nuclei are extracted and separately subjected to snHi-C. **(B**) Representative image of immunofluorescence staining of chromatin-bound MCM2 in wild type (WT) and in MCM-loading inhibited (MCM loss) G1 phase zygotes. Soluble proteins were pre-extracted. DNA is stained with DAPI. Scale bars: 10 μm. Quantification of mean chromatin-bound MCM2 intensity is depicted in fig. S2B and is based on 7 WT and 7 MCM loss zygotes. **(C)** Schematic representation of loop extrusion in as seen in single nucleus Hi-C (snHi-C) compared to bulk Hi-C. Extruding loops are stochastic leading to variable contacts in both bulk Hi-C and snHi-C maps. Convergently-oriented CTCF boundary elements can stall loop extrusion, resulting in anchored loops, which are detected in snHi-C upon averaging over the loop coordinates detected in MEFs. Changes in local chromatin compaction are studied by the contact probability, P_c_(s) as a function of genomic separation, s. **(D)** The strength of average loops and TADs for wild type (WT) and MCM-loading inhibited (MCM loss) shown separately for maternal and paternal nuclei. Data shown are based on n(WT, maternal) = 13, n(WT, paternal) = 16, n(No MCM, maternal) = 16, n(No MCM, paternal) = 15, from 4 independent experiments using 4-6 females. The heat maps were normalized to an equal number of reads. **(E)** P_c_(s) for maternal and paternal chromatin for control (WT) and MCM-loading inhibition (MCM loss) conditions. **(F)** Insulation score at TAD borders. The scores are calculated with a sliding diamond of 40 kb, with “0” position at the TAD boundary identified previously in CH12-LX cells.

Using this approach, we generated control and MCM loss zygotes, isolated maternal and paternal pronuclei in G1 phase and performed single-nucleus Hi-C (snHi-C) (Fig. 1A). Due to the sparsity of zygotic Hi-C data, it was not possible to perform *de novo* calling of loops (also referred as “corner peaks” or “dots”) that constitute local peaks of contact frequency and represent contacts between CTCF-bound loci. Instead, we used 12,000 loop coordinates from mouse embryonic fibroblast (MEF) Hi-C data that report on cohesin-dependent chromatin contacts in oocytes and zygotes (Fig. 1C) (31). Remarkably, MCM loss resulted in an increase in aggregate peaks and aggregate TADs (hereafter referred to simply as ‘peaks’ and ‘TADs’) in maternal chromatin and even a stronger increase in paternal zygotic chromatin (Fig. 1D). Paternal peaks were enriched for contact frequencies over intermediate (100-250 kb) and long-range interactions (>250 kb). Maternal peaks were more enriched in contact frequencies over longer distances (fig. S2C). The overall increase in aggregated Hi-C peak strengths upon MCM loss could reflect increased access of cohesins to CTCF sites, which is either directly due to changes in the cohesin loop extrusion process (potentially caused by loss of a barrier) or indirectly due to gene expression changes. To exclude the latter, we performed RNA-Seq and found no gross transcriptomic differences between control and MCM loss zygotes (fig. S3, A, B and C), which is consistent with the transcriptionally inactive state of these cells. We conclude that MCMs hinder the formation of CTCF-anchored loops and TADs independently of transcriptional changes.

To determine whether cohesins were responsible for the observed increase in Hi-C peak strength caused by MCM loss, we used a conditional genetic knockout approach based on Cre recombinase under control of the *Zp3* promoter to delete floxed alleles of the cohesin subunit *Scc1* in growing oocytes (2, 32). We microinjected mRNA encoding geminin^L26A^ in *Scc1^ΔΔ^* oocytes isolated from *ScclfW (Tg)Zp3-Cre* females and performed IVF to generate maternal *Scc1* knockout zygotes [*Scc1^Δ(m)/+(p)^*] (fig. S4A). Peaks and TADs were undetectable in *Scc1^Δ(m)+(p)^* zygotes, as reported previously (2), and remained undetectable if MCM loading is inhibited (fig. S4B). We conclude that MCMs interfere with cohesin-dependent loops and TADs.

To determine how MCMs affect chromatin organization, we examined genome-wide contact probability P_c_(s) curves as a function of genomic distance (s) (Fig. 1E and fig. S2F). Earlier we demonstrated that the shape of the P_c_(s) curve is informative of the extent of cohesin loop extrusion, i.e. the size of the shoulder in the curve reflects sizes of extruded loops (estimated using the derivative of log (P_c_(s)) (fig. S2G)) (2,33). Unexpectedly, MCM loss has little or no effect on the P_c_(s) curve of zygotic chromatin, suggesting that the process of loop extrusion is largely unaffected by MCM. This lack of change in the P_c_(s) curve contrasts the loss of the shoulder upon cohesin loss and the increase in the shoulder in *Wapl* knockout which considerably extends cohesin residence time (fig. S2E).

The overall lack of change in the contact probability P_c_(s) curve accompanied by changes in the “peak” intensities upon MCM loss is reminiscent of the effect of loss of CTCF (2,4,15). The effect of CTCF loss on “peaks”, however, is exactly opposite to that of MCM loss. We reason, and demonstrate below, that the presence of MCM partially interferes with the ability of CTCF to establish “peaks”, and MCM loss leads to strengthening of these CTCF-mediated features. If MCM impedes the establishment of CTCF-mediated structures, then MCM loss should increase CTCF peak strengths, as observed (Fig. 1D), which in turn may also lead to increased TAD boundary insulation. Indeed, without MCM, insulation increased for maternal and paternal zygotic chromatin (Fig. 1F). Together, these characteristic effects on chromatin organization are consistent with the notion that MCMs impede loop extrusion in a specific way, affecting largely CTCF-mediated peaks while having a small effect on sizes of extruded loops.

We therefore tested whether CTCF and MCM together determine the strength of TAD boundary insulation. We expressed geminin^L26A^ in CTCF knockdown oocytes isolated from *(Tg)Zp3-CTCFdsRNA* females and generated maternal CTCF knockdown zygotes (Fig. 2A and fig. S5A) (34). Remarkably, CTCF knockdown without or with MCM loss resulted in no detectable loops and TADs (Fig. 2B and fig. S5B). This effect was more severe than what has been observed by CTCF degradation in cell culture (4,15), suggesting efficient CTCF depletion during the OZT. There was no gross change in contact probability P_c_(s) curves, implying that extruding loops and chromatin compaction are unperturbed (fig. S5, C and D). CTCF knockdown caused weakening of TAD boundary insulation, which decreased further upon MCM loss, implying a barrier function for both CTCF and MCM (Fig. 2C). We conclude that despite the weak TAD structures in early mouse embryos, CTCF is essential for restricting CTCF-anchored loops and generating TAD boundaries within hours after fertilization. The lack of TAD organization in CTCF knockdown irrespective of MCM further suggests that, unlike CTCF, MCM has no instructive function for establishing boundaries, i.e. it does not stop cohesin at specific genomic locations. This is consistent with MCM barriers being located in different positions in different cells.

**Fig. 2.**
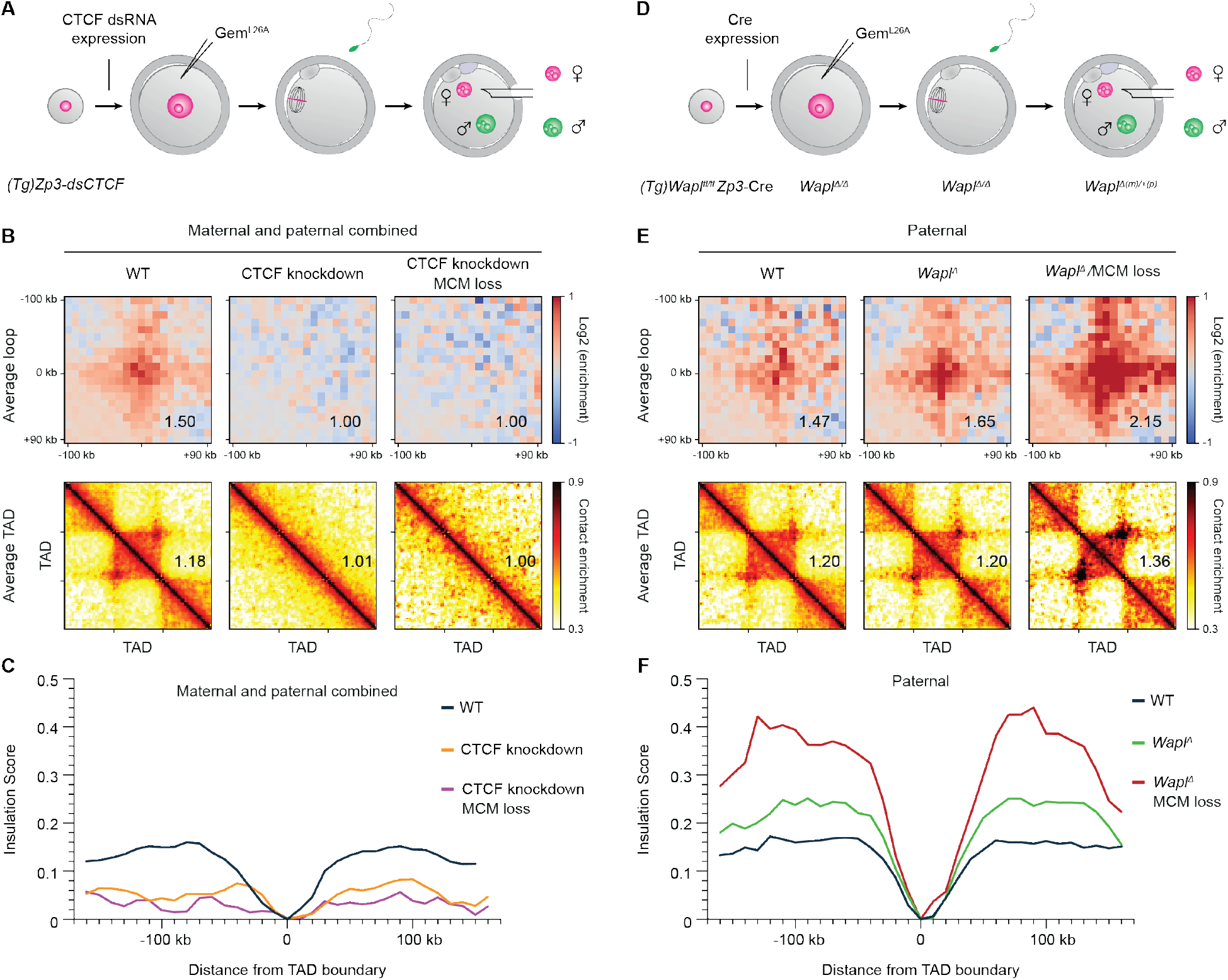
MCM hinders CTCF-anchored loops and functions independently of Wapl-mediated release of cohesin. (A) The *Zp3* promoter drives the expression of CTCF dsRNA in growing oocytes. CTCF depletion persists in mature (GV) oocytes and G1 zygotes. **(B)** The strength of average loops and TADs for wild type (WT), CTCF depleted (CTCF loss) and CTCF depleted combined with prevention of MCM loading (CTCF loss/MCM loss) in G1 zygotes. Maternal and paternal data are shown pooled together. Data are based on n(WT) = 21, n(CTCF loss) = 19, and n(CTCF loss/MCM loss) = 10 nuclei, from 3 independent experiments using 3 to 4 females for each genotype. The heat maps were normalized to an equal number of reads. **(C)** Insulation score at TAD borders from combined maternal and paternal nuclei. Data shown are based on same number of samples as in aggregate loop and TAD analysis. **(D)** Generation of conditional genetic maternal knockout zygotes based on *(Tg)Zp3*-Cre. The *Zp3* promoter initiates expression of Cre recombinase during oocyte growth which deletes the conditional (floxed) alleles of *Wapl*. After fertilization, maternal knockout zygotes *Wapl^Δ(m)/+(p)^* are generated. Because most proteins are provided by the oocyte and the G1 phase zygote is practically transcriptionally inactive, these maternal knockout zygotes are depleted for Wapl. **(E)** The strength of average loops and TADs for control (WT), Wapl depleted (*WaplΔ*) and Wapl depleted combined with prevention of MCM-loading (*Wapl^Δ^/MCM* loss) for paternal nuclei in G1 zygotes. Data shown are based on n(*Wapl^fl^*,paternal) = 18, n(*Wapl^Δ^/MCM*, paternal) = 8, n(*Wapl^Δ^/MCM* loss, paternal) = 5. Control samples are WT samples (this study) pooled with *Wapl^fl^* samples (previously published in (2)). The heat maps were normalized to an equal number of reads. **(F)** Insulation score at TAD borders for paternal nuclei. Data shown are based on same number of samples as in aggregate loop and TAD analysis.

We also considered an alternative possibility that MCMs affect cohesin loops by functioning with Wapl in releasing cohesin from chromatin. This hypothesis is based on the similarity of the effect on Hi-C peak and TAD strengths observed following *Wapl* knockout and MCM loss (2,4,35), where the strongest effects occurred for MCM loss in paternal chromatin (Fig. 2E). If Wapl and MCMs function together, then their co-depletion is expected to resemble the individual depletions. Alternatively, if they function independently, then co-depletion could have synergistic effects. To distinguish between these, we expressed geminin^L26A^ in *Wapl^Δ/Δ^* oocytes isolated from *Wapl^fl/fl^(Tg)Zp3-Cre* females and generated maternal *Wapl* knockout zygotes [*Wapl^Δ(m)/+(p)^*] (Fig. 2D) (31). Combined MCM loss and *Wapl* knockout strongly increased peak and TAD strengths over the separate conditions (Fig. 2E and fig. S6A), suggesting that they function independently. The combined loss also had strong synergistic effects on increasing TAD boundary insulation, suggesting that MCMs still restrict loop extrusion when cohesin residence time is increased (Fig. 2F and fig. S6B). These results provide further evidence for a barrier effect of MCMs on zygotic chromatin organization.

Since these findings were obtained in a specialized system, we tested their relevance for human somatic cells. Transient transfection of geminin^L26A^ into synchronized HCT116 cells only mildly reduced MCM abundance on chromatin (data not shown), precluding this approach for further analysis. To directly degrade MCMs, we treated synchronized HCT116 cells carrying endogenously tagged auxin-inducible degron MCM2-(mAID) alleles with DMSO or auxin (fig. S7, A and B) (36). Auxin treatment from mitosis until G1 phase reduced MCM2 and MCM4 abundance without grossly affecting cohesin abundance (fig. S7C). We generated bulk Hi-C maps and observed small enrichments in corner peaks on Hi-C maps and in aggregate peak and TAD analysis for MCM-depleted versus control cells (fig. S7, D, E, F and G). These effects are consistent with the findings in zygotes but less pronounced; this could be due to more efficient depletion of loaded MCMs during the OZT and/or MCMs having a more dominant barrier effect on zygotic “ground state” chromatin, which might have fewer other DNA binding proteins. We conclude that MCMs also impede the formation of CTCF-anchored loops and TADs in somatic cells.

We further tested whether MCM loading affects gene expression. Acute MCM degradation resulted in differential expression of 454 transcripts (FDR = 0.1) (fig. S8, A, B and C). The magnitude of the gene expression effects is roughly comparable to loss of cohesin, CTCF and Wapl (3,15,35). When analysing Hi-C contact frequencies around transcription start sites (TSS), we found little change for non-differentially expressed transcripts and an increase in contact frequencies for both up- and down-regulated transcripts between control and MCM2-depleted conditions (fig. S8D). The promoters of genes whose expression changed >1.5-fold in either direction showed the most pronounced change in contact frequencies, implying that these are somehow sensitive to changes in chromatin organization. These effects could be indirect, but it is tempting to speculate that loop extrusion dynamics affected by obstacles like MCMs alter gene expression.

The most parsimonious interpretation of the effect of MCM loss on Hi-C data is that MCMs interfere with cohesin loop extrusion by forming randomly located extrusion barriers. If so, then we predicted that MCMs are physical obstacles to cohesin on DNA. To directly test this, we established an MCM roadblock assay for passive cohesin translocation using total internal reflection fluorescence (TIRF) microscopy that detects real-time cohesin-MCM interactions at the single-molecule level. To this end, origin licensing was reconstituted from purified components in a stepwise manner on origin-containing DNA molecules, 21 kb in length, tethered at both ends to a slide surface (Fig. 3A). Robust loading of MCM and double-hexamer formation, a hallmark of proper origin licensing, was observed in the presence of the initiation factors ORC, Cdc6 and Cdt1 (37). Cohesin was subsequently introduced in moderate salt conditions promoting translocation on fast timescales (fig. S9B and movie S1) (38).

**Fig. 3.**
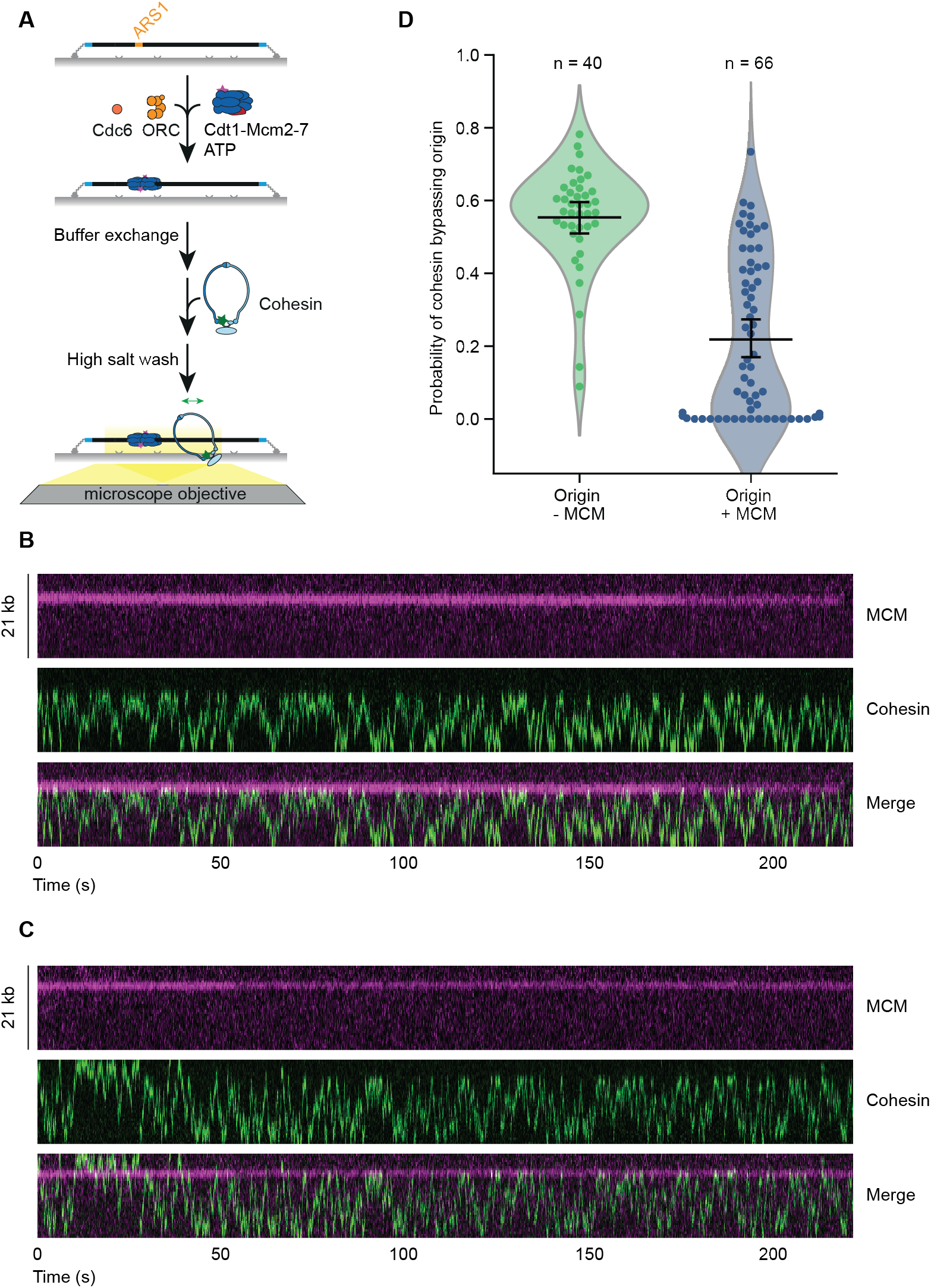
MCM is a barrier for cohesin translocation *in vitro.* (**A**) Schematic principle of a singlemolecule cohesin translocation assay on licensed DNA. Initially MCM is loaded onto DNA in presence of the licensing factors ORC and Cdc6, followed by cohesin. Subsequently, cohesin translocation is visualized in absence of free protein and buffer flow. (**B-C**) Representative kymographs of translocating cohesin on licensed DNA. Origin-bound MCM is an efficient barrier for cohesin translocation (**B**) but still supports occasional passage (**C**) during a 220 s observation interval. (**D**) MCM is a barrier for cohesin translocation. Probability of translocating cohesin bypassing the origin in absence or presence of MCM calculated from 40 or 66 molecules with 8025 or 10732 visualized encounters, respectively. Black lines display the mean within a 95 % confidence interval (generated by bootstrapping). See Material and Methods section for details.

Direct visualisation of cohesin encounters with the potential roadblock revealed that MCMs frequently constrained cohesin translocation (Fig. 3, B and C, fig. S9, C and D, and movies S2 and S3). To quantify origin passing probability, we used subpixel localisation and tracking and found that passage is three times less frequent when MCMs are present (Fig. 3D). This difference was independent of the cohesin diffusion coefficient, which remained unchanged in the presence of MCMs (Fig. S9E). We found that MCMs are strong barriers (defined as an origin passing probability of <10%) for ~50% of cohesin molecules (31/66, Fig. 3D). Since some cohesin molecules were unable to pass origins even once during the 220 second imaging window, despite frequent attempts (15/66, Fig. 3B and fig. S9B), MCMs can also be an impermeable barrier over these timescales. The failure to pass the origin was specific to MCMs, as all cohesin molecules readily translocated over origins without MCMs (40/40, fig. S9B). These origins likely also lacked ORC because the complex dissociates in the high salt wash. To exclude the possibility that multiple MCM complexes were loaded, which might generate an artificial obstacle, we performed photobleaching studies to distinguish between single, double and multiple hexamers, and found that 65% of MCMs were loaded as double hexamers (fig. S9A). Interestingly, single MCM hexamers were sufficient to confine cohesin on one side of the origin (4/7, fig. S9F), raising the possibility that single hexamers could also impede loop extrusion and active MCMs may displace cohesin ahead of the replication fork. This speculation assumes that an obstacle to diffusive cohesin translocation will also impair active loop extrusion. However, a limitation of this assay is that we do not know if passive translocation and loop extrusion are related. It will be important to test whether functionally loaded MCMs halt active loop extrusion, which requires a combined assay that has so far not been possible to establish due to both reactions occurring under specific conditions *in vitro.* We conclude that MCMs are physical barriers to cohesin translocation, which may occasionally be bypassed.

To consolidate the *in vivo* and *in vitro* data, and to test the hypothesis about the mechanism of MCM action, we created a quantitative model of MCMs as randomly located barriers to cohesin loop extrusion and performed polymer simulations. In our model (Fig. 4A), cohesins extrude loops bidirectionally by bringing together DNA flanking their loading point. Extrusion of a loop is stopped (in one direction) if cohesins encounter: 1) another cohesin, 2) a CTCF site (~ 50% of the time as estimated experimentally), or 3) an MCM complex (~ 20% of the time, as identified by a parameter sweep - see below) (fig. S10, S11, S12 and S13). CTCF and MCM can both stall loop extrusion but may allow extruding cohesin to bypass these barriers some of the time, consistent with observations from the single-molecule experiments. Strikingly, we found that it is possible to observe increased peak strength at CTCF sites (Fig. 4B) as observed experimentally (Fig. 1D), without strongly affecting contact probability curves, i.e. without affecting the mean size of extruded loops.

**Fig. 4.**
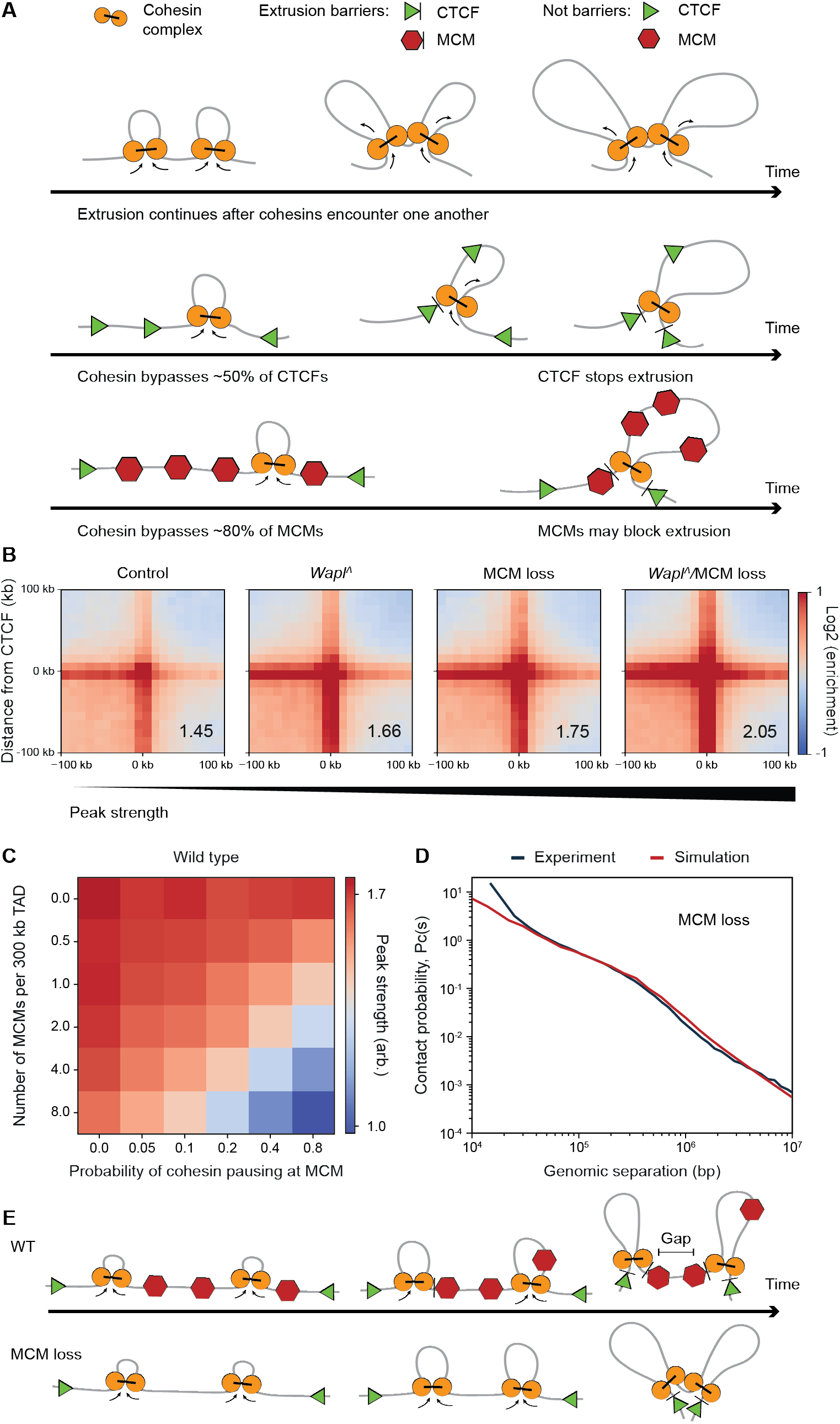
Simulation model of MCM as a barrier to cohesin loop extrusion. (**A**) Schematic illustration of a quantitative model for loop extrusion by cohesin and interactions with CTCF and MCM complexes. Cohesin complexes (yellow) extrude loops. On encountering barriers such as CTCF sites (green triangles), MCM complexes (red hexagons), or other cohesins, extrusion can be blocked. Cohesins may bypass some CTCFs and MCMs, but not others; the choice to bypass an MCM or CTCF site is stochastic and varies in time. **(B-D)** For the paternal chromatin simulations, we assume CTCFs stall cohesins ~45% of the time as measured in mESCs (40) and MCMs stall cohesins 20% of the time. **(B**) Peak strengths for simulated paternal chromatin under various genetic perturbations for TADs of average length 300 kb. **(C)** Matrix of peak strengths showing a linear trade-off between the MCM density and its ability to pause cohesins. **(D)** Simulated contact probability decay curve P_c_(s) for MCM loss condition is well matched with the experimental data. **(E)** Model summarizing our finding that chromatin-bound MCMs can function as barrier for loop extrusion in G1 phase.

To obtain the best fitting model described above, we performed a sweep of simulation parameters, such as cohesin processivity and linear density, as well as parameters of MCM density and permeability. Simulations allowed us to identify a narrow range of parameters for each condition (WT, MCM loss, *Wapl* knockout and double MCM loss/*Wapl* knockout) such that both the peak strengths and P_c_(s) curves of paternal chromatin can be quantitatively reproduced (Fig. 4, C and D, fig. S10, S11, S12, S13 and S14). The simulations suggest that in WT and MCM loss conditions, cohesins extrude paternal chromatin loops of 110-130 kb and have a density of ~1/(300 kb). Interestingly, to achieve the simultaneous increase in peak strength without strongly affecting the average cohesin loop length after MCM loss, it was necessary for MCMs to be permeable to cohesin loop extrusion; there was a linear trade-off between the MCM density and its ability to pause cohesins (Fig. 4C). We found that for experimentally estimated densities of MCMs between 1 MCM/(30 kb) to 1 MCM/(150 kb), cohesins must bypass MCMs in about 60-90% of encounters. Thus, for an estimated density (see Supplementary Materials) of 1 MCM/(75 kb), cohesins will pause at an MCM for ~20% of encounters (Fig. 4C). Simulations independently provide similar values of MCM density and permeability when the effect of MCM loss in *Wapl* knockout is reproduced. This permeability value is larger than what we observed by single-molecule experiments but could reflect the different conditions, i.e. *in vivo,* on chromatin, and with loop extrusion, and could explain how CTCF-anchored loops are generated in the presence of MCMs in G1 phase cells. Thus, our simulations provide further support for the model that MCMs are semi-permeable barriers to loop extrusion *in vivo*.

Our models also provide a rationale for this seemingly contradictory ability of MCM to reduce the strength of CTCF-mediated features without affecting sizes of extruded loops. CTCF-mediated peaks of contact probability emerge if one or more cohesins extrude loops in a region between neighbouring CTCF barriers. Such peaks, however, get considerably weaker or disappear if a gap is present between extruded loops (39). On the contrary, sizes of extruded loops are unaffected by such gaps between consecutive loops. If one or more randomly located barriers are present between a pair of CTCFs, extrusion is likely to leave some unextruded gaps that result in weaker CTCF-CTCF peaks (Fig. 4E). If separation between such random barriers is large (i.e. comparable to average cohesin loop size of ~100kb) or the barriers are sufficiently permeable, then the barriers have little effect on sizes of extruded loops. This is new and a rather surprising effect of random barriers on different features of chromosome organization.

In summary, we have identified MCM complexes as barriers for loop extrusion based on *in vivo*, *in vitro* and simulation data. MCMs are members of a new class of randomly positioned, abundant and cell cycle phase-specific obstacles that impede CTCF-anchored loop and TAD formation. Which features, beyond size or specific interaction motifs (such as the case of CTCF), determine whether a macromolecular complex impedes the loop extrusion process remain to be determined.

It is conceivable that the topological association of proteins with DNA plays an important role in modulating loop extrusion and hence chromosome organization. Consistent with this, cohesin complexes that topologically entrap sister chromatids are likely barriers to other cohesin complexes mediating loop extrusion. It is also interesting to consider whether MCMs might be ancestral barriers in species lacking CTCF-anchored loops such as *Drosophila*, where TAD establishment during embryonic development coincides with a switch in replication origin usage. Lastly, our data suggest that the “MCM paradox” has unexpected consequences for chromatin organization and potentially gene expression, which might have relevance for human pathologies, such as Meier-Gorlin syndrome, that are linked to mutations in the MCM loading pathway.

## Acknowledgements

We are grateful for generous contributions of the *(Tg)Zp3-dsCTCF* mouse strain from M.S. Bartolomei and the *MCM2-mAID* HCT116 cell line from M. Kanemaki. We thank the Remus lab for generating MCM strains used for *in vitro* single-molecule assays. We are grateful to Z. Lygerou for sharing geminin^L26A^-GFP plasmid. We thank H.C. Theußl and J.R. Wojciechowski (IMP/IMBA Transgenics facility) for reviving *(Tg)Zp3-dsCTCF* embryos. Illumina sequencing was performed at VBCF NGS Unit and MPIB NGS core facility (http://www.vbcf.ac. at and https://www.biochem.mpg.de/7076201/NGS). **Funding:** B.J.H.D. is funded by the PhD program (DK) Chromosome Dynamics (W1238-B20), supported by the Austrian Science Fund (FWF). L.A.M. and H.B.B. acknowledge support from the National Institutes of Health Common Fund 4D Nucleome Program (DK107980) and the Human Frontier Science Program (HFSP RGP0057/2018). I.F. is supported by Medical Research Council UK University Unit grant (MC_UU_00007/2). Research in the Duderstadt laboratory is supported by the European Research Council (ERC-StG-804098 ReplisomeBypass), the German Research Foundation (DFG, SFB863-111166240), and the Max Planck Society. Research in the J.-M.P. laboratory is supported by Boehringer Ingelheim, Austrian Research Promotion Agency (Headquarter grant FFG-852936), ERC (Grant Agreement No 693949), HFSP (RGP0057/2018) and Vienna Science and Technology Fund (grant LS19-029). Research in the Tachibana laboratory is supported by the ERC (ERC-CoG-818556 TotipotentZygotChrom), HFSP (RGP0057/2018), Herzfelder foundation and FWF (P30613-B21), Austrian Academy of Sciences and Max Planck Society. **Author contributions:** K.T. conceived the project. B.J.H.D. supervised by K.T. performed snHi-C, bulk Hi-C, RNA-Seq and imaging experiments. H.B.B. and M.J.S contributed equally to the project. H.B.B. supervised by L.A.M. developed and performed snHi-C data analysis and polymer simulations. M.S.J. supervised by K.E.D. performed single-molecule imaging and analysis. J.G. performed snHi-C and imaging experiments. S.P. performed snHi-C data analysis and RNA-Seq analysis. I.G. performed RNA-Seq analysis. I.M.F. performed bulk Hi-C analysis. I.F.D. supervised by J.-M.P. purified recombinant cohesin^STAG1, SCC1^-^Halo^. B.J.H.D., H.B.B., M.J.S, I.M.F. and I.G. prepared the figures. B.J.H.D., H.B.B., M.J.S, I.G., K.E.D., L.A.M. and K.T. wrote the manuscript with input from all authors.

---

**Fig. S1.**
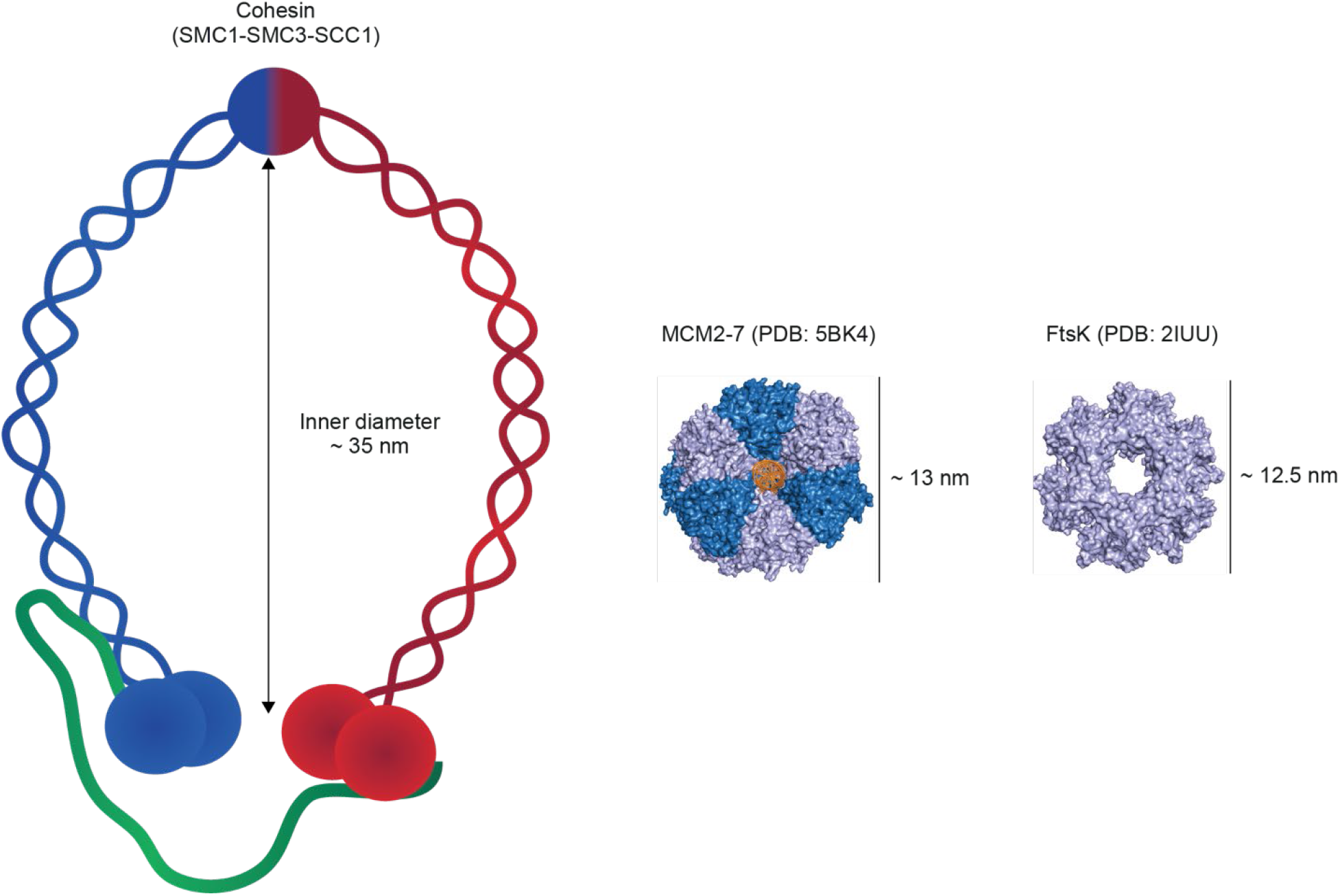
Dimension of cohesin compared to the estimated size of the MCM2-7 complex and the FtsK roadblock. Size comparison between a schematic representation of the ring-shaped heterotrimeric cohesin (SMC1-SMC3-SCC1) with the MCM2-7 double hexamer and the FtsK monohexamer. The size estimation of MCM2-7 and FtsK was based on their crystal structure. PDB accessions codes for each protein are shown in parentheses.

**Fig. S2.**
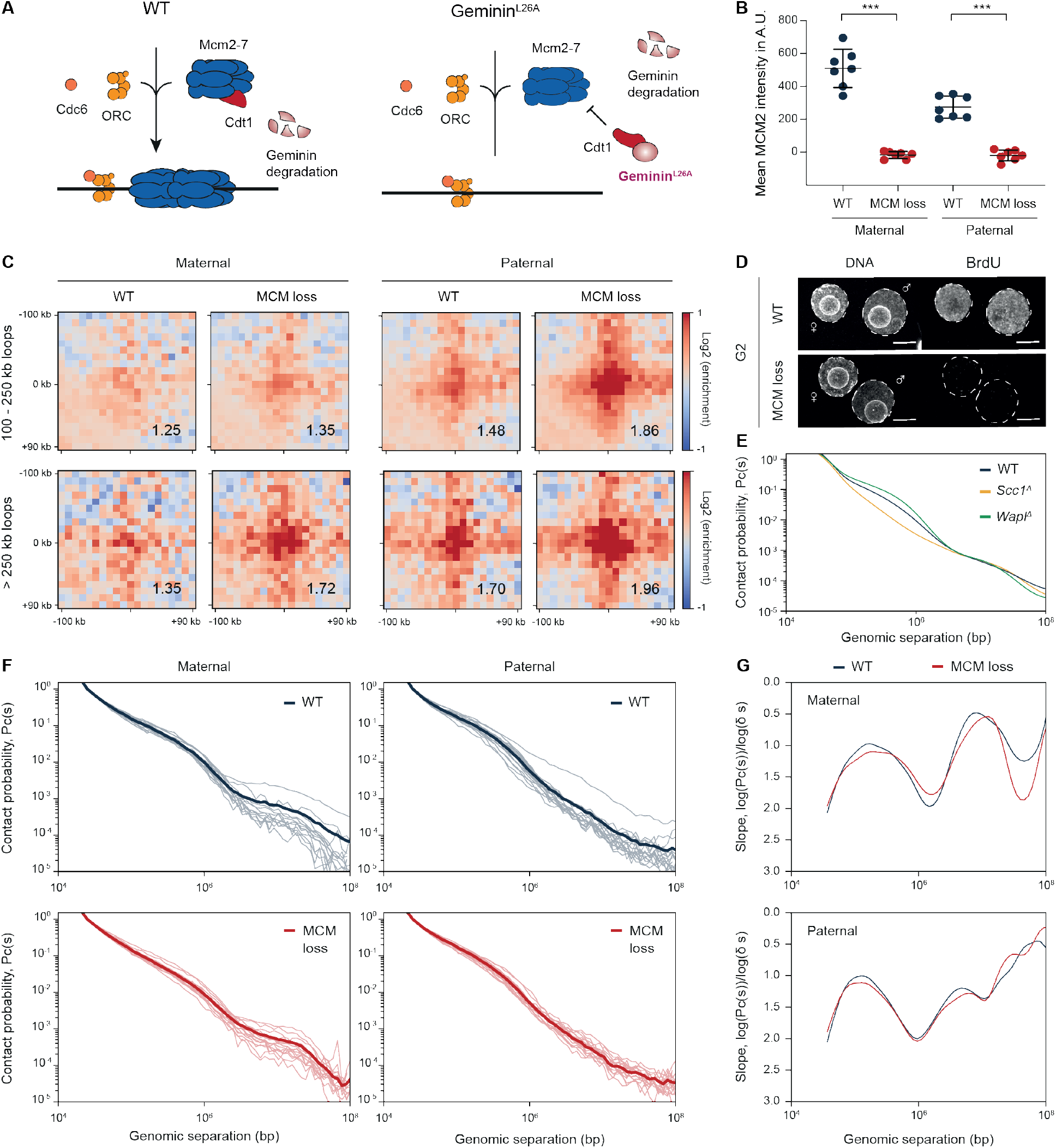
Inhibition of MCM loading increases loop strength in G1 zygotes without affecting local chromatin compaction. **(A)** Expression of geminin^L26A^, a non-degradable variant of geminin, prevents the recruitment of MCMs to chromatin in G1 phase by inhibiting the Cdt1-mediated loading pathway. **(B)** Quantification of mean chromatin-bound MCM2 intensity in wild type (WT) and MCM-loading inhibited (MCM loss) G1 phase zygotes. Soluble proteins were preextracted. *** p <0.001, calculated using unpaired t-test. Representative image is shown in Fig. 1B. **(C)** Average aggregate loop strength separated in intermediate (100-250kb) and long loops (>250kb) for wildtype (WT) and MCM-loading inhibited (MCM loss) conditions for maternal and paternal nuclei. Data shown are based on same number of samples as in Fig. 1D. **(D)** Representative image for immunofluorescence analysis to detect BrdU incorporation in G2 phase cells from 2 WT and 4 geminin^L26A^-injected cells. The cells were cultured in continuous presence of BrdU since 4 h post-fertilization until fixation. DNA is stained with DAPI. Scale bars: 10 μm. **(E)** Contact probability P_c_(s) curve as a function of genomic distance (s). Cohesin is directly involved in shaping the P_c_(s) in the range 0.3-3 Mb, which represent local chromatin compaction. The contact frequency in this region decreases upon cohesin depletion (*Scc1^Δ^*) but increases upon enrichment of chromatin-bound cohesin (*Wapl^Δ^*) **(F)** Contact probability P_c_(s) curves for individual G1 zygotes with average P_c_(s) (same as in Fig. 1E) in bold overlayed. **(G)** Slopes of the P_c_(s) curves (depicted in Fig. 1E) as an indication for the average size of cohesin-extruded loops.

**Fig. S3.**
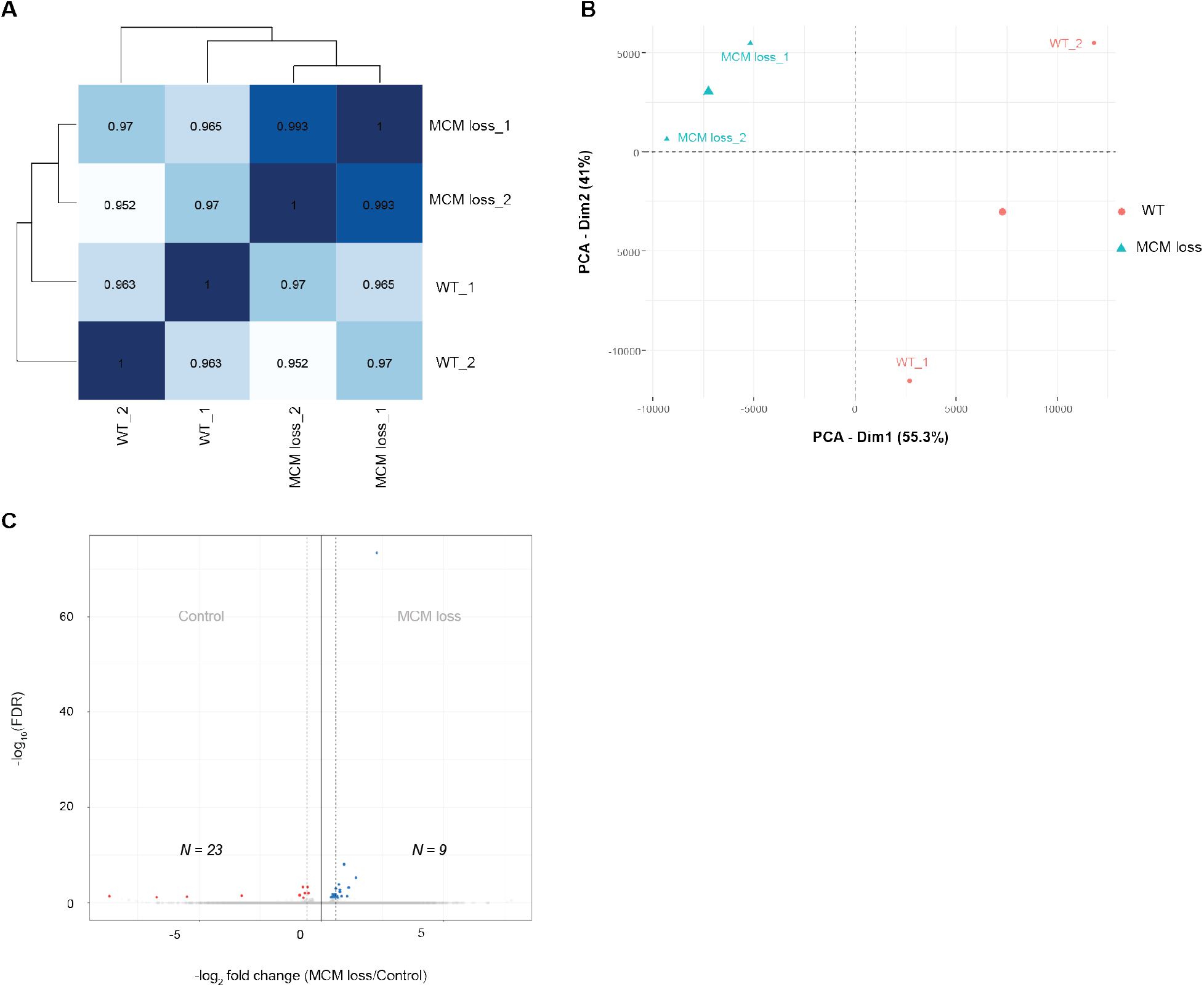
Gene expression in G1 zygotes is largely unaffected when MCM recruitment is prevented. **(A)** Heatmap of Euclidean distances between the transcriptomes of wild type control (WT_1 & 2) and MCM-loading inhibited (MCM loss_1 & 2) G1 zygotes. Numbers indicate pairwise Pearson’s correlation r. **(B)** Principal component analysis of the transcriptomes of wild type control (WT_1&2) and MCM-loading inhibited (MCM loss_1&2) G1 zygotes. **(C)** Volcano plot showing statistical significance –log^10^ (FDR) versus fold change (log^2^ fold change) for RNA-seq data between wild type (WT control) and MCM loading-inhibited (MCM loss) conditions. Numbers indicate the number of transcripts significantly up-(right) or downregulated (left) upon MCM loss at FDR=0.1. Dashed vertical lines indicate −0.585 and +0.585 log_2_ fold change in expression (1.5-fold decrease and increase in expression), respectively.

**Fig. S4.**
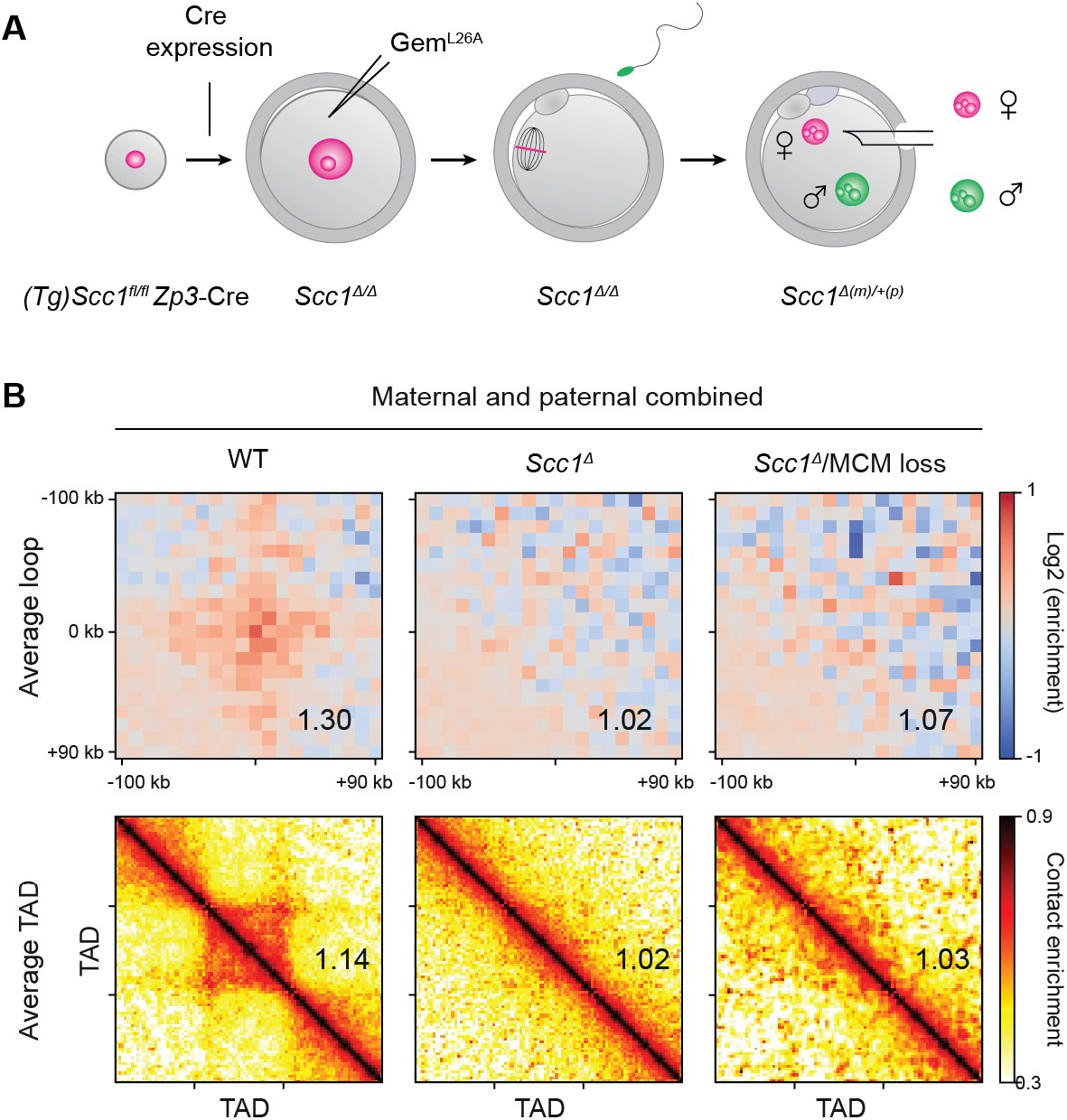
MCM impedes cohesin-dependent loops and TADs. **(A)** Generation of conditional genetic maternal *Scc1* knockout zygotes based on *(Tg)Zp3-Cre.* The *Zp3* promoter initiates expression of Cre recombinase during oocyte growth which deletes the conditional (floxed) alleles of *Scc1.* After fertilization, maternal knockout zygotes Scc1 ^*Δ(m)/+(p)*^ are generated. **(B)** The strength of average loops and TADs for control (*Scc1^fl^*), Scc1 depletion (Scc1^Δ^) and Scc1 depletion with prevention of MCM-loading (*Scc1^Δ^/MCM* loss). Maternal and paternal data are shown pooled together. Data are based on n(*Scc1^fl^*) = 26, n(*Scc1^Δ^*) = 44, and n(*Scc1^Δ^/MCM* loss) = 10 nuclei. The heat maps were normalized to an equal number of reads. Control (*Scc1^fl^*) samples and 38 Scc1-depleted samples (*Scc1^Δ^*) were previously published (2).

**Fig. S5.**
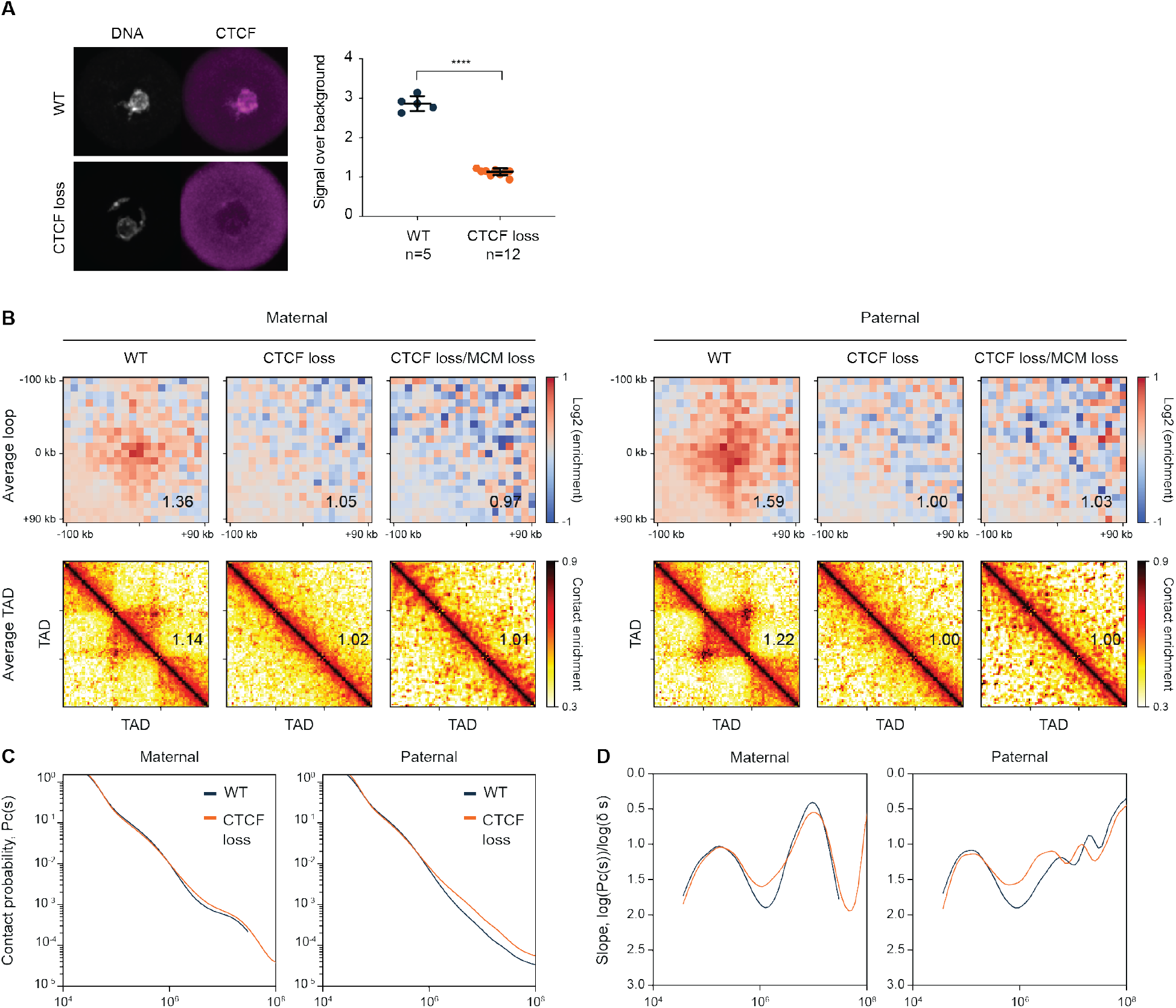
CTCF is required for loop and TAD formation in G1 zygotes. **(A)** Immunofluorescence analysis of CTCF in wild-type and CTCF-depleted GV oocytes. (Left) Representative images: DNA is stained with DAPI. (Right) Quantification of CTCF intensity. **(B)** The strength of average loops and TADs for wild type (WT), CTCF depleted (CTCF loss) and CTCF depleted combined with prevention of MCM loading (CTCF loss/MCM loss) in G1 zygotes, shown separately for maternal and paternal nuclei. Data shown are based on n(WT, maternal) = 10, n(WT, paternal) = 11, n(CTCF loss, maternal) = 11, n(CTCF loss, paternal) = 8, n(CTCF loss/MCM loss, maternal) = 6, n(CTCF loss/MCM loss, paternal) = 4 nuclei. The heat maps were normalized to an equal number of reads. **(C)** Contact probability P_c_(s) curve for wild type (WT) and CTCF depleted (CTCF loss) conditions, shown separately for maternal and paternal nuclei. **(D)** Slopes of the P_c_(s) curves (panel C) as an indication for the average size of cohesin-extruded loops.

**Fig. S6.**
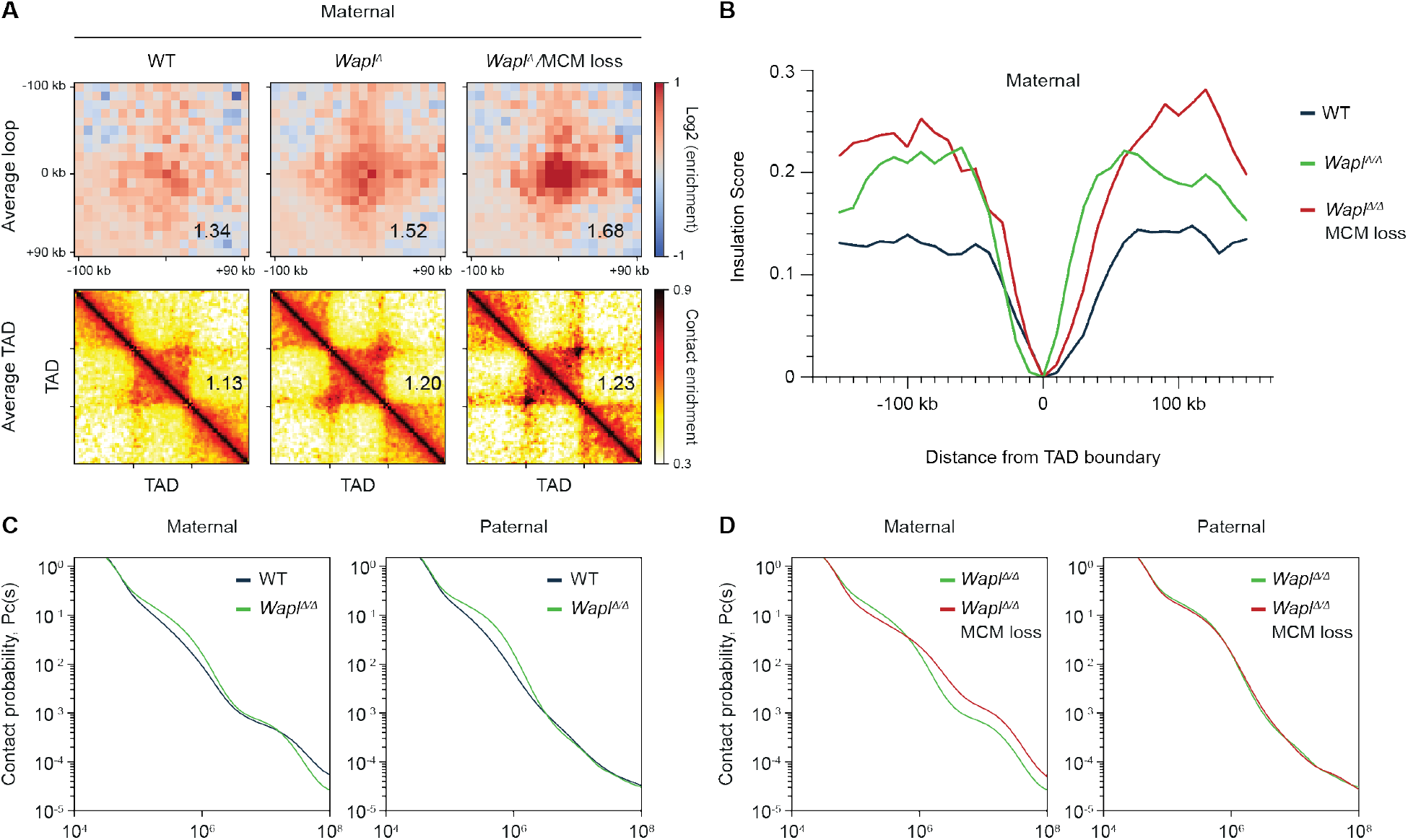
MCM restricts loops and TAD formation in maternal chromatin independently of Wapl-mediated cohesin release. **(A)** The strength of average loops and TADs for control (WT), Wapl depleted (*Wapl^Δ^*) and Wapl depleted combined with prevention of MCM loading (*Wαpl^Δ^/MCM* loss) for maternal nuclei in G1 zygotes. Data shown are based on n(*Wapl^fl^*, maternal) = 18, n(*Wαpl^Δ^*, maternal) = 8, n(*Wαpl^Δ^*/MCM loss, maternal) = 7. Control samples are WT samples (this study) pooled with *Wapl^fl^* samples (previously published in (2)). **(B)** Insulation score at TAD borders for maternal nuclei. Data shown are based on same number of samples as in aggregate loop and TAD analysis. **(C-D)** Contact probability P_c_(s) curves for control (WT), Wapl depleted (*Wapl^Δ^*), and Wapl depleted combined with prevention of MCM-loading (*Wapl^Δ^*/MCM loss) conditions, shown separately for maternal and paternal nuclei.

**Fig. S7.**
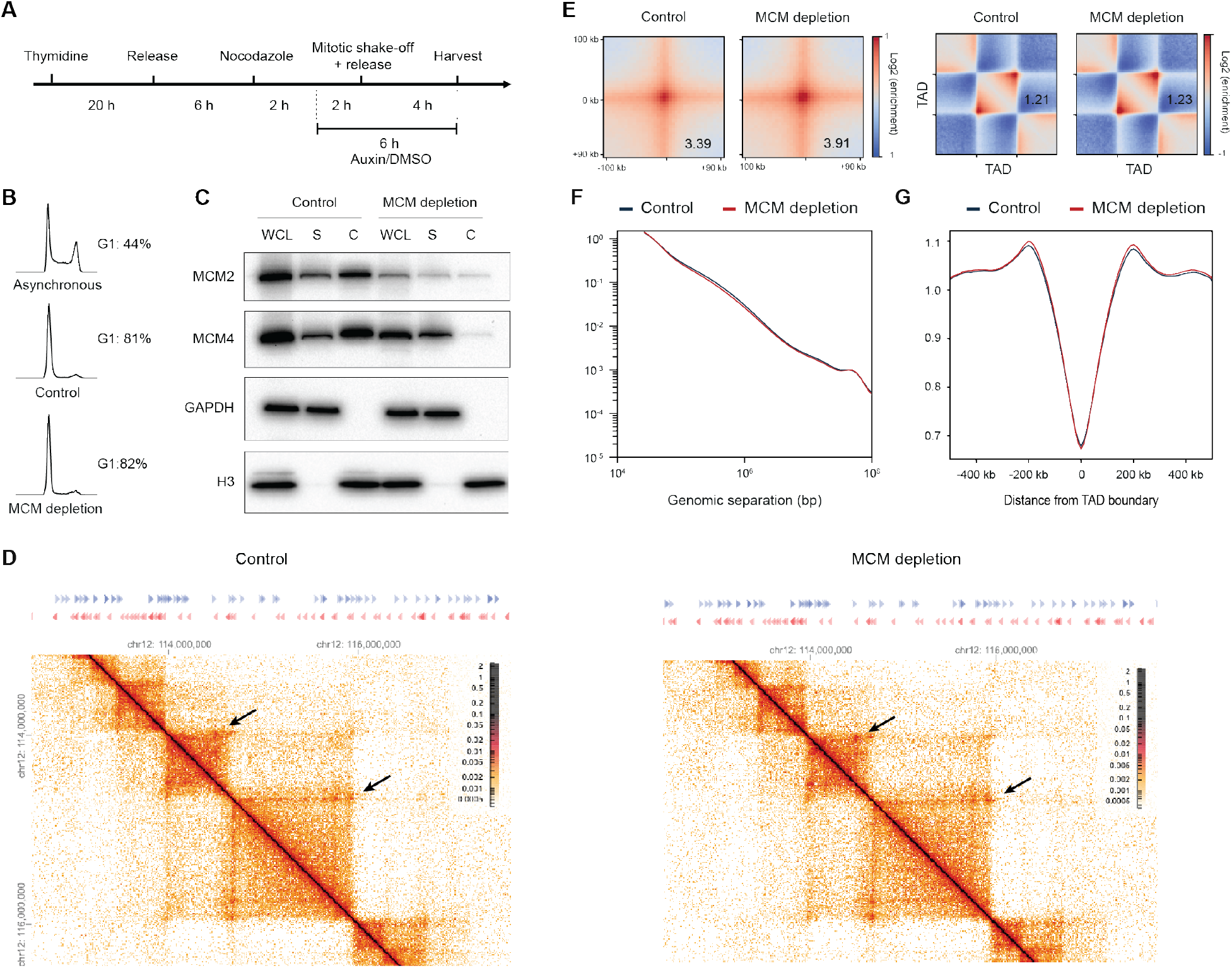
Acute MCM2 depletion results in an increase of average loops and TADs and leads to a mild increase in TAD insulation. (**A**) Schematic for G1 synchronization of HCT116 MCM2-mAID cells. (**B**) Cell cycle profiles of HCT116 MCM2-mAID cells (Control/MCM loss) synchronized in G1 compared to asynchronous cells. (**C**) Immunoblotting analysis of whole-cell lysate (WCL), supernatant fraction (S) and chromatin fraction (C) from HCT116 MCM2-AID treated with DMSO (Control) or auxin (MCM loss) for MCM2, GAPDH and H3. GAPDH and H3 are used as a control for the supernatant and chromatin fraction, respectively. (**D**) Hi-C contact matrices from HCT116 MCM2-AID treated with DMSO (Control) or auxin (MCM loss) for the region 112,5-117,6 Mb on chromosome 12 at 10 kb resolution. Increased corner peaks are denoted with an arrow. CTCF sites are depicted above the contact matrices. (**E**) Average of the total contact frequency of loops and TADs in an aggregate peak analysis for control and MCM loss cells. (**F**) Contact probability P_c_(s) curves for DMSO (Control) and auxin treatment (MCM loss). (**G**) Insulation pileups for DMSO (Control) and auxin treatment (MCM loss). The ‘insulation strength’ is the average enrichment of the top-left and bottom-right corners, divided by the average enrichment of the top-right and bottom-left corners (central transition bins are not included).

**Fig. S8.**
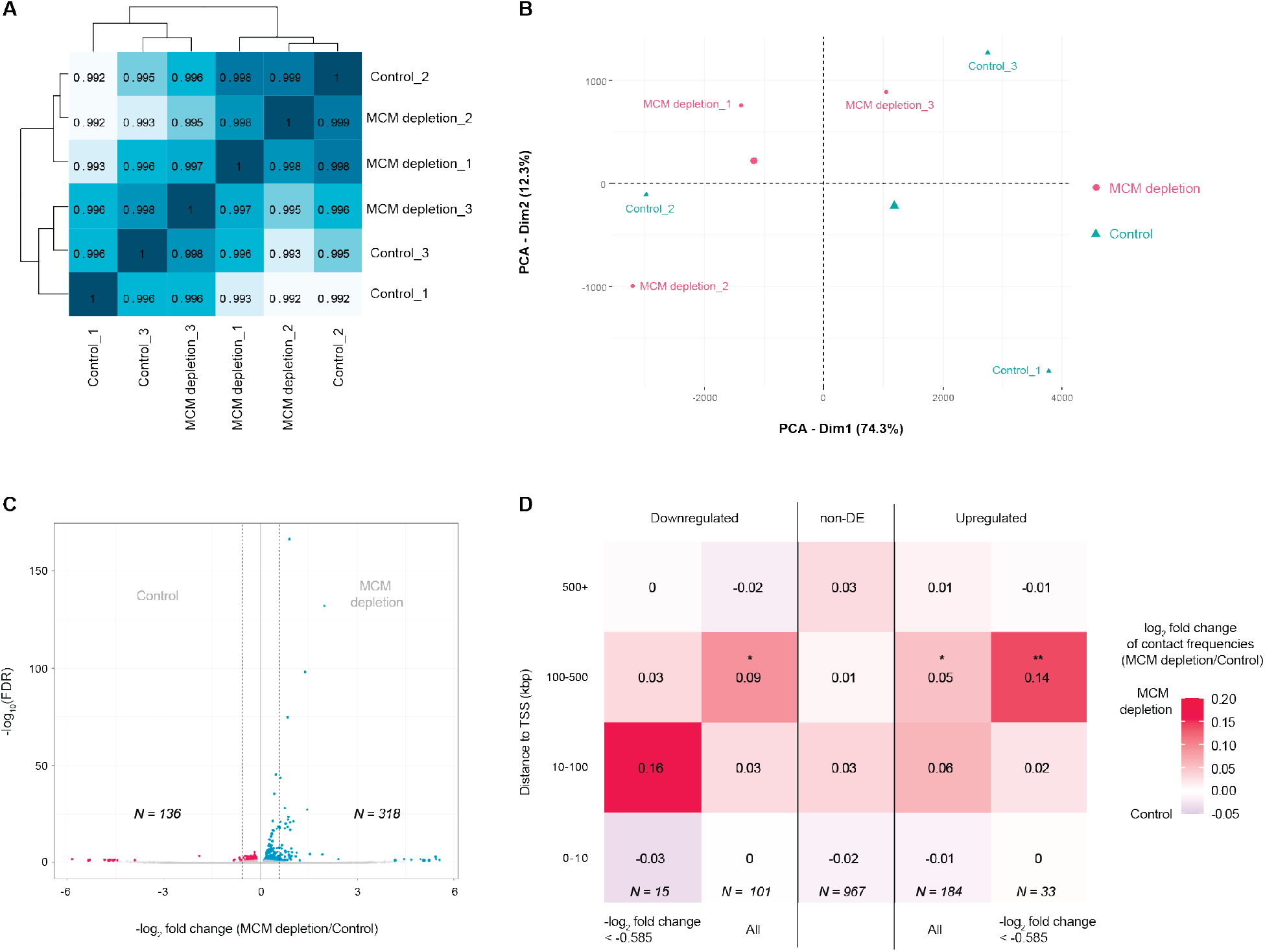
Differential gene expression upon MCM depletion in somatic cells. (**A**) Heatmap of Euclidean distances between the transcriptomes of MCM-AID expressing HCT116 cell treated with auxin (MCM depletion_1, 2 & 3) or DMSO (Control_1, 2 & 3). Numbers indicate pairwise Pearson’s correlation r. (**B**) Principal component analysis of the transcriptomes of MCM-AID expressing HCT116 cell treated with auxin (MCM depletion_1, 2 & 3) or DMSO (Control_1, 2 & 3). (**C**) Volcano plot showing statistical significance –log^10^ (FDR) versus fold change (log^2^ fold change) for RNA-seq data between MCM-AID expressing HCT116 cells treated with DMSO or auxin. Numbers indicate the number of transcripts significantly up-(right) or downregulated (left) upon MCM loss at FDR=0.1. Dashed vertical lines indicate −0.585 and +0.585 log_2_ fold change in expression (1.5-fold decrease and increase in expression), respectively. (**D**) Heatmap of mean contact frequency changes of the TSS of differentially expressed transcripts (DE TSS) between DMSO or auxin treated MCM-AID expressing HCT116 cells. Rows indicate the mean change of contact frequency in the indicated distance range from the TSS. Middle (3^rd^) column indicates mean contact frequency changes of non-differentially expressed TSS used as control (non-DE TSS). Columns 2 and 4 shows mean contact frequency changes of down- and upregulated TSS, respectively, columns 1 and 5 show the same data for TSS whose transcripts underwent at least 1.5-fold (log_2_ −0.585) decrease or 1.5-fold increase (log_2_ 0.585), respectively. Numbers in the center of cells show the log_2_ of mean contact frequency change. * and ** indicates significant difference (p<0.05 and p< 0.01, respectively) in the mean contact frequency change between the DE TSS and the corresponding bin of non-DE TSS. Numbers in italics indicate the number of TSS used for analyzing the corresponding column. Note that the discrepancy of numbers to panel (**C**) is either due to multiple TSS sharing the same bin in the Hi-C analysis or the corresponding bin containing the TSS missing from the Hi-C analysis.

**Fig. S9.**
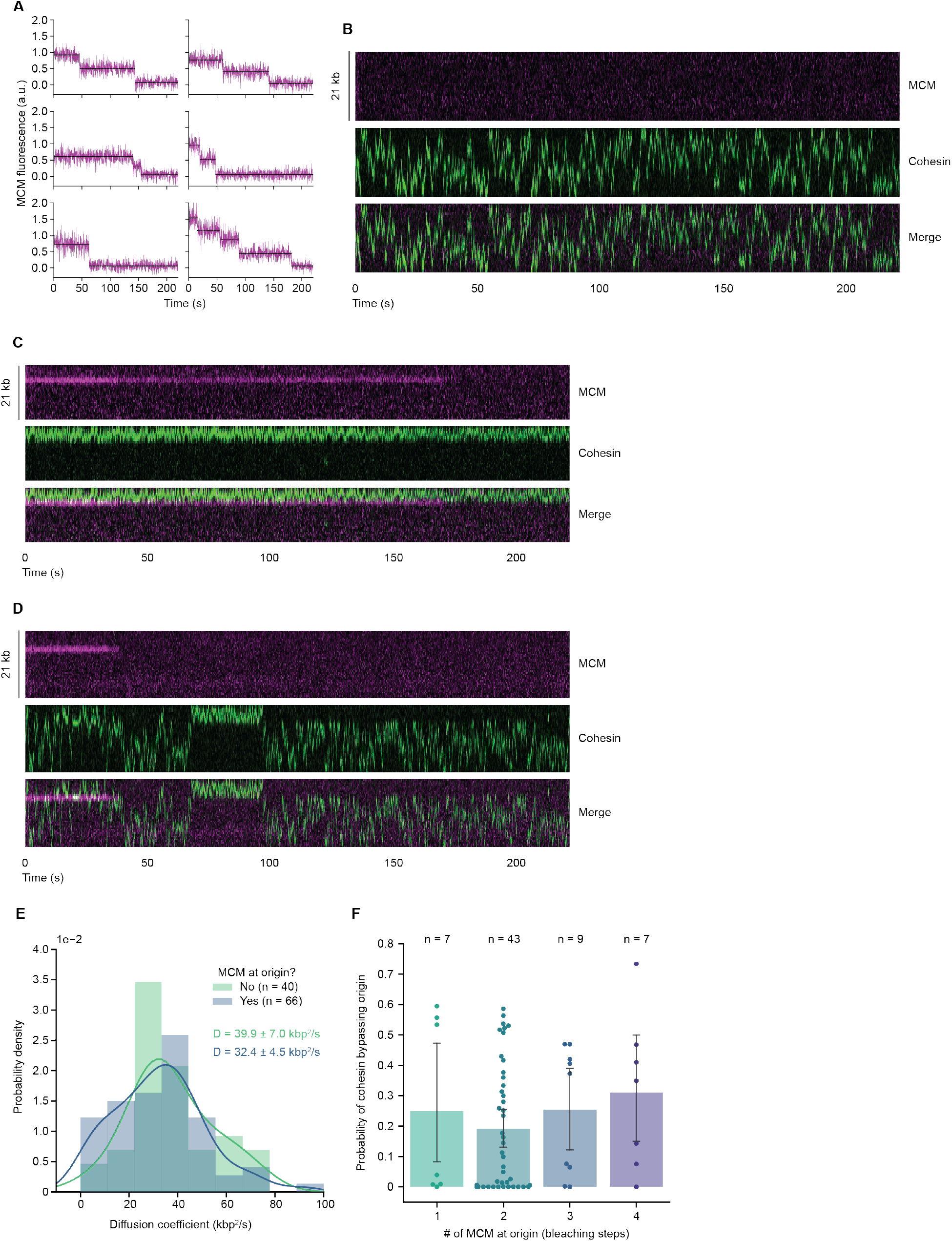
Translocating cohesin can bypass MCM with reduced efficiency. (**A**) MCM loads as double-hexamers at origins. Time traces of origin bound MCM fluorescence intensity (purple) show two-step bleaching but less frequent also one and multi-step (black, fits by kinetic change point analysis) (**B-D**) Representative kymographs of translocating cohesin in absence (**B**) or presence (**C** and **D**) of MCM at the origin. MCM is an efficient barrier for cohesin translocation (C) but intervals of efficient and inefficient MCM passage are observed (**D**) during a 220 s interval. All kymographs were generated along a line based on subpixel fitting of the ends of individual DNAs. DNAs doubly-tethered with different extension and as a consequence, kymographs differ in heights. These length differences were accounted for throughout all analysis steps. (**E**) MCM does not alter observed cohesin translocation velocity. Distribution of diffusion coefficients (D) of translocating cohesin in presence or absence of MCM. Lines display the kernel density estimation and mean +/−1.96 × SEM is shown. (**F**) Multiple loaded MCM do not increase the barrier strength for cohesin translocation. Photobleaching analysis confirm loaded MCM double-hexamers as main species. Error bars (95 % confidence interval) were generated by bootstrapping.

**Fig. S10.**
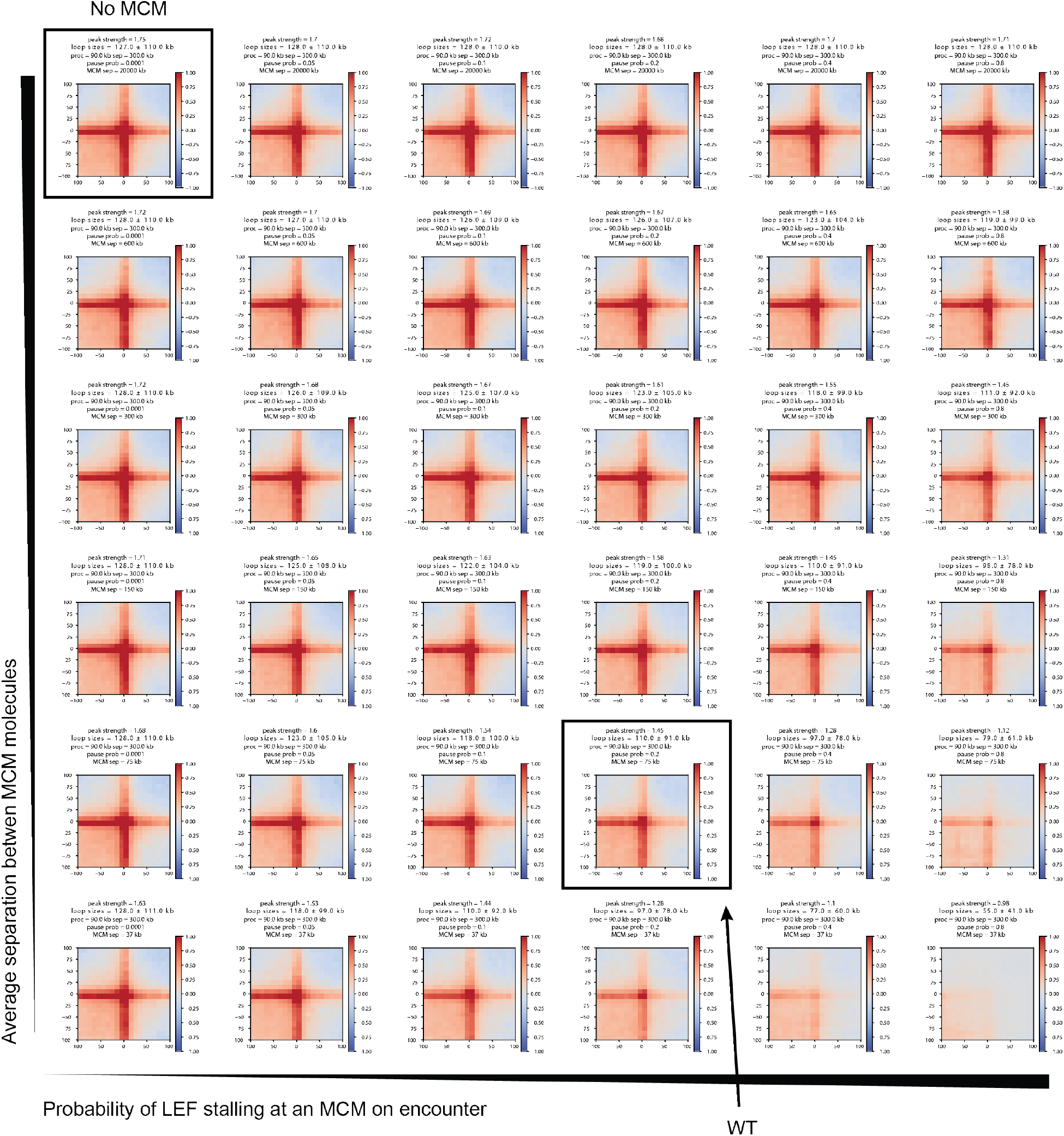
Parameter sweep for peak strengths for simulated paternal chromatin in control and MCM loss conditions.

**Fig. S11.**
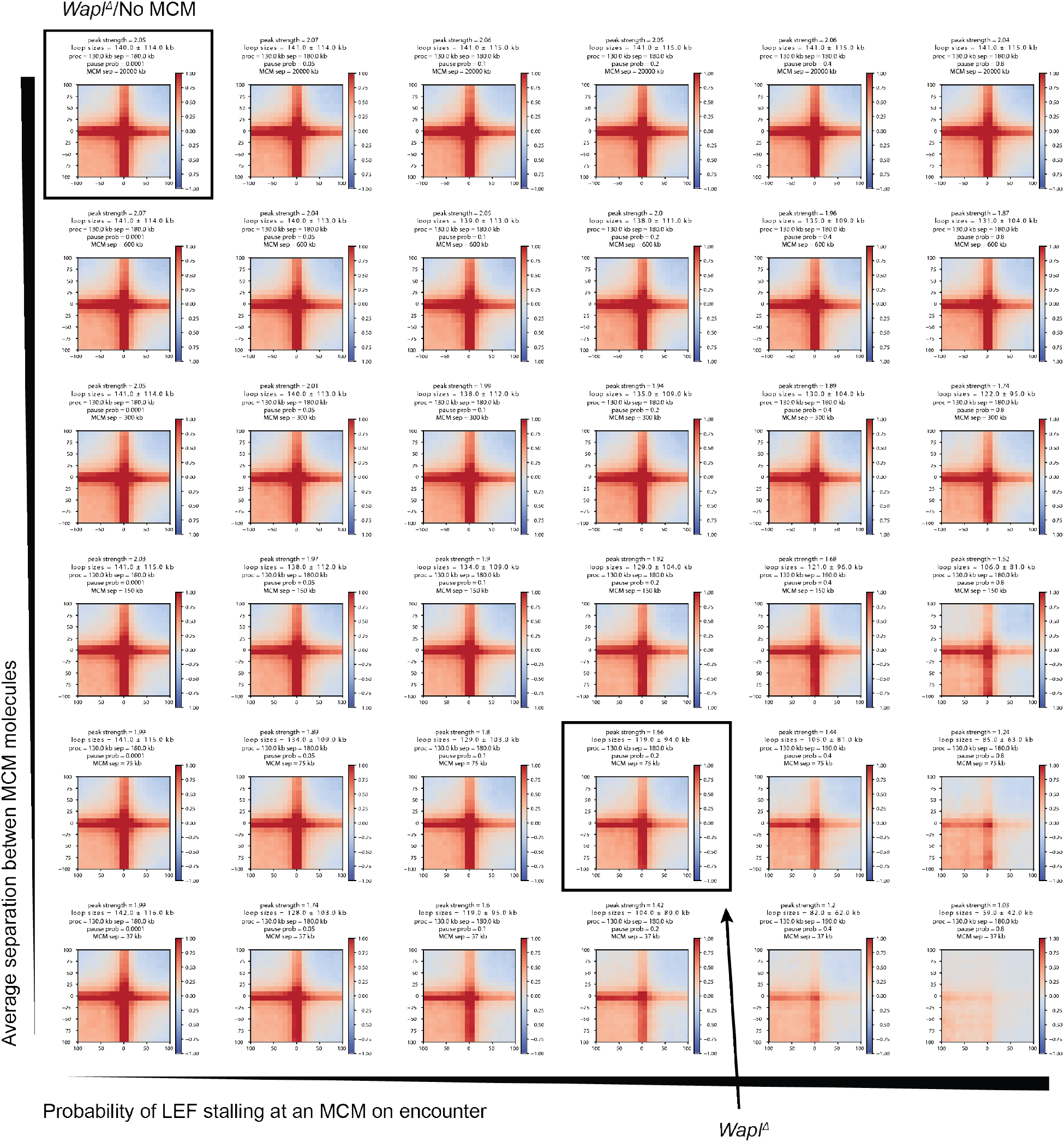
Parameter sweep for peak strengths for simulated paternal chromatin in *Wapl^∆^* and *Wapl^∆^*/MCM loss conditions.

**Fig. S12.**
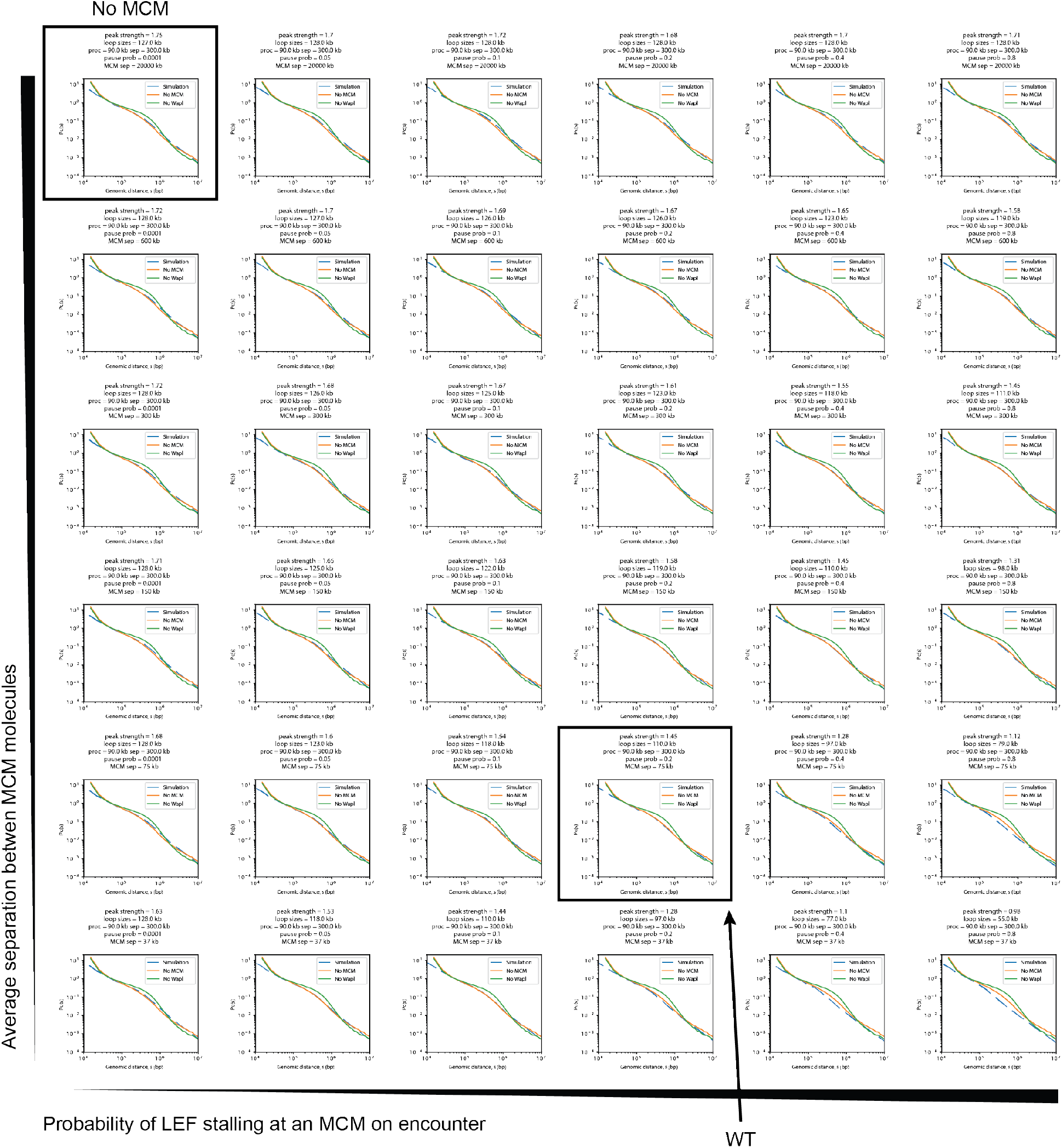
Parameter sweep for contact probability decay curves P_c_(s) for simulated paternal chromatin in control and MCM loss conditions.

**Fig. S13.**
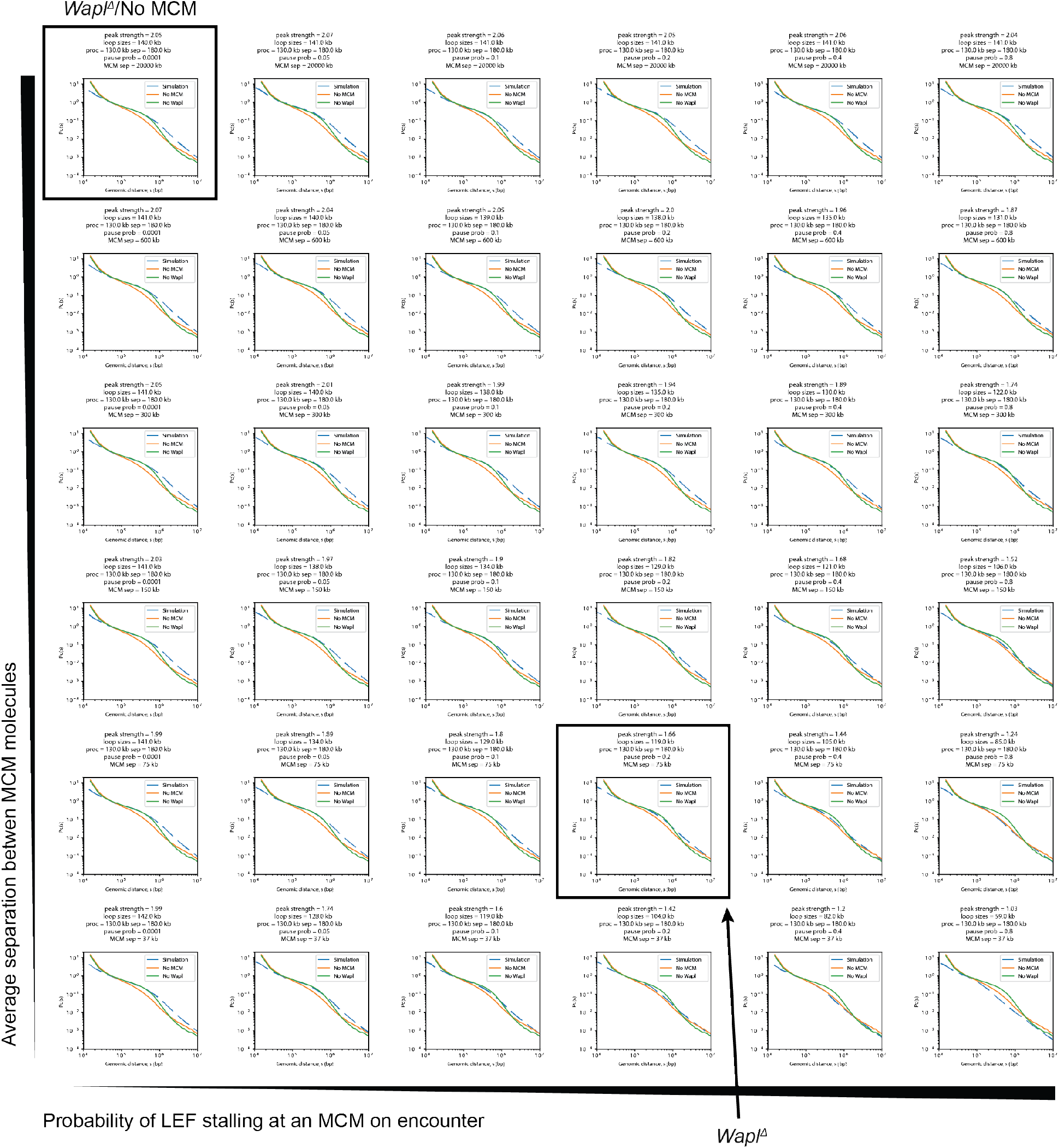
Parameter sweep for contact probability decay curves P_c_(s) for simulated paternal chromatin in *Wapl^∆^* and *Wapl^∆^*/MCM loss conditions.

**Fig. S14.**
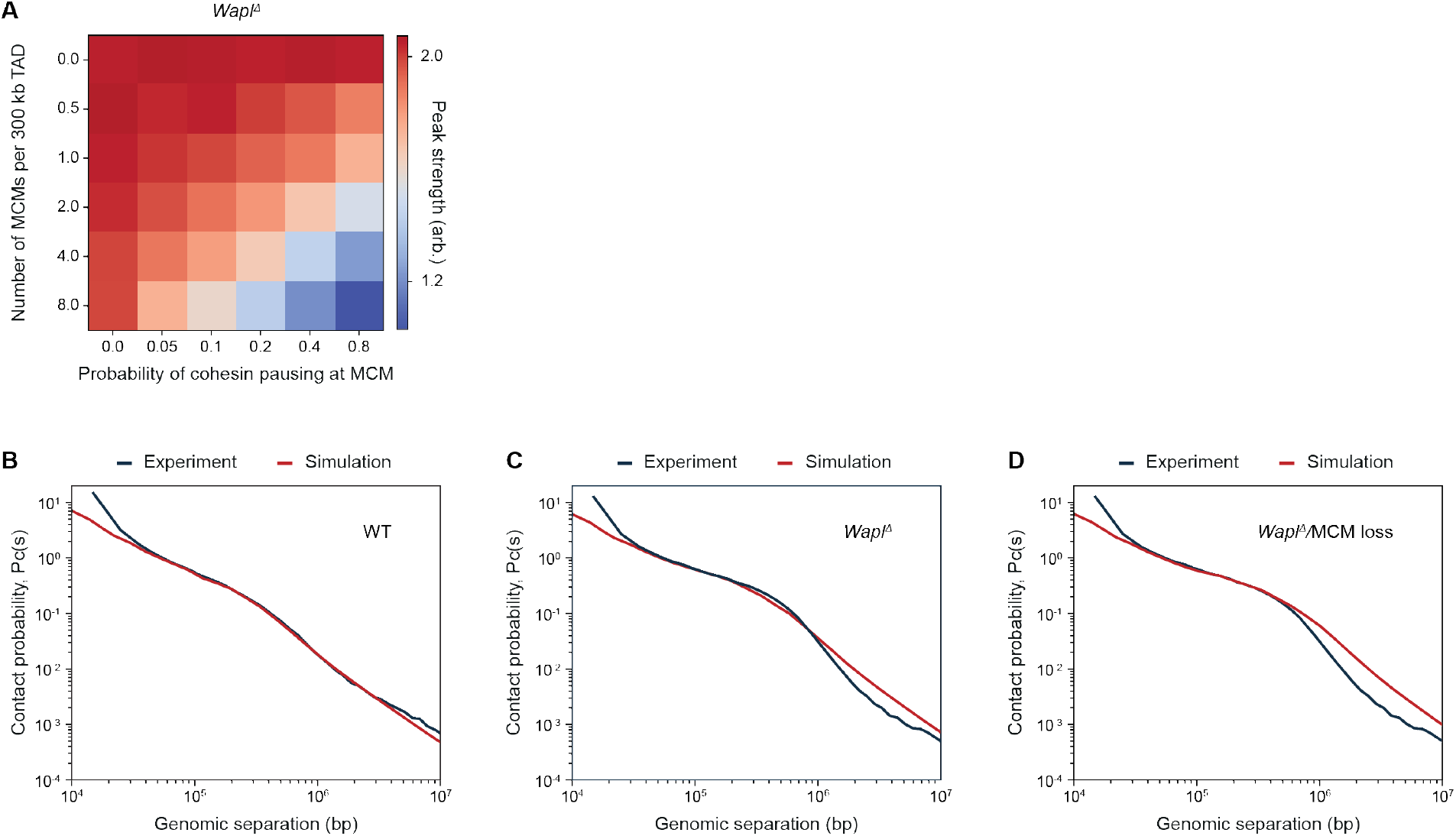
Polymer simulations of MCM as a barrier to cohesin loop extrusion. (**A**) Matrix of peak strengths showing a linear trade-off between the MCM density and its ability to pause cohesins in *Wapl^∆^.***(B-D)** Simulated contact probability decay curve P_c_(s) for WT **(B)**, *Wapl^∆^***(C)** and *Wapl^∆^*/MCM loss **(D)**. The simulated P_c_(s) curve is well matched with the experimental data.

## Captions for Movies S1 to S3

**Movie S1.** Movie showing translocating cohesin (green) on doubly-tethered DNA (blue) in absence of MCM (corresponds to kymograph in fig. S9B).

**Movie S2.** Movie showing origin-bound MCM (magenta) as an efficient barrier for cohesin translocation (green) on doubly-tethered DNA (blue) (corresponds to kymograph in Fig. 3B).

**Movie S3.** Movie showing translocating cohesin (green) on doubly-tethered DNA (blue) that occasionally can bypass origin-bound MCM (magenta) (corresponds to kymograph in Fig. 3C).

## Materials and Methods

### Animals

The mice used in this work were bred and maintained in agreement with the authorizing committee according to the Austrian Animal Welfare law and the guidelines of the International Guiding Principles for Biomedical Research Involving Animals (CIOMS, the Council for International Organizations of Medical Sciences). Mice were housed in individually ventilated cages at a daily cycle of 14-h and 10-h dark with access to food and water *at libitum.* All mice were bred in the IMBA animal facility. Wildtype and *Sccl^fl/fl^* mice were bred on a mixed background (B6, 129, Sv). *Wapl^fl/fl^* and *Zp3-dsCTCF* mice were bred on a primarily C57BL/6J background. *Zp3-dsCTCF* mice were maintained by breeding *Zp3-dsCTCF* males to C57BL/6J females. Experimental *Sccl^fl/fl^* and *Wapl^fl/fl^* mice were obtained by mating of homozygous floxed females to homozygous floxed males carrying *Tg(Zp3Cre)* (4ĩ, 42). Experimental *Zp3-dsCTCF* mice were maintained by breeding *Zp3-dsCTCF* males to C57BL/6J females.

### Mouse oocyte collection and *in vitro* culture

Ovaries were dissected from sexually mature female mice, which were sacrificed by cervical dislocation. Fully grown germinal vesicle (GV) oocytes from 2-5 months old females were isolated by physical disaggregation of ovaries with hypodermic needles. GV oocytes were cultured in M2 medium supplemented with 0.2 mM of the phosphodiesterase inhibitor 3-Isobutyl-1- methylxanthine (IBMX, Sigma-Aldrich) at 37 °C. Mature oocytes were selected according to appearance (size, central nucleus, smooth zona pellucida) and cultured in M16 medium supplemented with IBMX in an incubator at 37°C and 5% CO2. Oocytes were cultivated in ~40 μl drops covered with paraffin oil (NidOil).

### Microinjection

GV oocytes were microinjected with *in vitro* transcribed mRNA dissolved in RNAse-free water (mMessage mMachine T3 kit, Ambion). Following mRNA concentrations have been injected: 2.3 pmol hGeminin^L26A^ and 0.2 pmol GFP. Microinjection was performed in ~20 μl drops of M2 (0.2 mM IBMX) covered with mineral oil (Sigma-Aldrich) using a Pneumatic PicoPump (World Precision Instruments) and hydraulic micromanipulator (Narishige) mounted onto a Zeiss Axiovert 200 microscope equipped with a 10x/0.3 EC plan-neofluar and 40x/0.6 LD Apochromat objective. Injected oocytes were cultured for 2 h and then released from IBMX inhibition by washing in M16 to resume meiosis.

### *In vitro* maturation and *in vitro* fertilization

Oocyte collection and culturing was performed as described above but M2 and M16 media was supplemented with 20% FBS (Gibco) and 6 mg/ml fetuin (Sigma-Aldrich). After microinjection and IBMX release as described above, GV oocytes were subsequently incubated at 37°C and low oxygen conditions (5% CO_2_, 5% O_2_, 90% N_2_) to initiate *in vitro* maturation to metaphase II (MII) eggs. Next, MII eggs were *in vitro* fertilized 10.5-12h post-release of IBMX. Sperm was isolated from the *cauda epididymis* and *vas deferens* of stud males and capacitated in fertilization medium (Cook Austria GmbH) in a tilted cell culture dish for at least 30 min. before incubation with MII eggs. For *in vitro* fertilization of wildtype, *Sccl^f/fl^ Tg(Zp3-Cre)* and *Zp3-dsCTCF* oocytes, sperm was obtained from B6CBAF1 males, while sperm of C57BL/6J males was used for *in vitro* fertilization of *Wapl^fl/fl^ Tg(Zp3-Cre)* oocytes. Zygotes were scored by the formation of visible pronuclei at 5.5-7 h post-fertilization.

### *In situ* fixation, immunofluorescence staining and image acquisition

Zygotes were pulsed with 1 mM EdU or BrdU (Invitrogen) before *in situ* fixation to check the time frame of G1 phase. To check for DNA replication, zygotes were fixed in G2 after continuous incubation in presence of EdU or BrdU. Oocytes and zygotes were stripped from there zona pellucida by using acidic Tyrode’s solution (Sigma-Aldrich) before *in situ* fixation in 4% PFA (in PBS) for 30 min at RT, followed by permeabilization in 0.2% Triton X-100 in PBS (PBSTX) for 30 min at RT. EdU-pulsed cells were processed according to the manual of the Click-iT™ EdU Alexa Fluor 647 imaging kit (Invitrogen). Blocking was performed using 10% goat serum (Dako) in PBSTX for 1 h at RT or at 4°C overnight. Cells were incubated with primary antibodies for 2.5 h at RT or at 4°C overnight. Following primary antibodies were used: anti-MCM2 (1:500; BD transduction Laboratories, #610701), anti-CTCF (1:250, Peters laboratory A992). After washing in blocking solution three times for at least 20 min, cells were incubated with Alexa Fluor^®^ 488, 568 or 647 secondary antibodies (Invitrogen) for 1 h at RT. Excess of secondary antibody wash removed by washing three times in 0.2% PBSTX for at least 20 min, followed by short PBS wash and 20 min submerged in Vectashield^®^ with DAPI (Vector Labs). Cells were mounted in Vectashield^®^ with DAPI using imaging spacers (Sigma-Aldrich) to preserve 3D integrity.

Detection of chromatin-bound MCM2 required pre-extraction before fixation and was performed with minor modifications as described in (43). In short, zona pellucida was not removed and zygotes were incubated in ice-cold extraction buffer (50 mM NaCl; 3 mM MgCl2; 300 mM Sucrose; 25 mM HEPES; 0.5% Triton X-100) for 7 min on ice, followed by three short washes in ice-cold extraction buffer without Triton X-100. *In situ* fixation and immunofluorescence was performed as described above. To avoid zona pellucida collapse, cells were submerged in increasing Vectashield concentrations before final mounting.

Image acquisition was performed on a Zeiss LSM780 confocal microscope using a planapochromat 63x/1.4 oil immersion objective.

### Cell culture and synchronization

HCT116 cells were cultured as previously described (36). Briefly, cells were cultured in McCoy’s 5A medium (Thermo Fisher Scientific) supplemented with 10% FBS (Gibco), 2 mM L-glutamine (Invitrogen) and 10% penicillin-streptomycin solution (Sigma-Aldrich). Cells were grown in an incubator at 37°C with 5% CO2. MCM2-mAID degradation was induced by addition of 500 μM 3-indoleacetic acid (Sigma-Aldrich) for 6 h. To synchronize cells in G1, a 2 mM thymidine arrest was followed by release into fresh medium for 6 h. Subsequently, nocodazole was added for 5 h, followed by shake-off of prometaphase cells and release in fresh medium for 4 h. Cells were fixed for Hi-C, microscopy and FACS. Cell cycle profiling was performed using propidium iodide staining.

### Chromatin fractionation and protein detection

Fractionation was performed as previous described (4). Briefly, cells were extracted in a buffer consisting of 20 mM Tris-HCl (pH 7.5), 100 mM NaCl, 5 mM MgCl, 2 mM NaF, 10% glycerol, 0.2% NP40, 20 mM b-glycerophosphate, 0.5 mM DTT, and protease inhibitor cocktail (Complete EDTA-free, Roche). Chromatin pellets and supernatant were separated and collected by centrifugation at 1,700 g for 5 min. The chromatin pellets were washed three times with the same buffer. Protein concentration was measured using a Bradford assay. Proteins were separated through SDS-PAGE on a Bolt™ 4-12% Bis-Tris Plus Gel (Invitrogen) and transferred to a nitrocellulose membrane. After overnight blocking with 5% skim milk in TBS-T at 4°C, the membrane was incubated with primary antibodies for 2,5 h at RT. Following antibodies were used: anti-MCM2 (1:5000; BD transduction Laboratories, #610701), anti-MCM4 (1:5000; Abcam, #ab4459), anti-H3(1:2000; Cell Signaling, #97155), anti-GAPDH(1:2500; Millipore, #MAB374). Goat Anti-Mouse/Rabbit Immunoglobulins/HRP (Dako) secondary antibodies were used to detect primary antibodies. Detection was performed using Immobilon Forte Western HRP Substrate (Merck) with a ChemiDoc imaging system (Bio-Rad).

### Single nucleus Hi-C

Single nucleus Hi-C was carried out as previously described (2, 28, 31, 44). Pronuclei of wildtype, *Scc1^∆/∆^, Wapl ^∆/∆^* and *Zp3-dsCTCF* zygotes were fixed around 1.5 hours post-visualization of pronuclei (corresponding to 6-7.5 h post-fertilization) and therefore are expected to be in G1 phase of the cell cycle. No blinding or randomization was used for handling of the cells. Briefly, isolated pronuclei were fixed in 2% PFA for 15 min, transferred to microwell plates (Sigma, M0815) and then lysed on ice in lysis buffer (10 mM Tris-HCl pH 8.0; 10 mM NaCl; 0.5% (v/v) NP-40 substitute (Sigma); 1% (v/v) Triton X-100 (Sigma); 1x Halt™ Protease Inhibitor Cocktail (Thermo Scientific)) for at least 30 min. After a brief PBS wash, the pronuclei were incubated in 1x NEB3 buffer (New England Biolabs) with 0.6% SDS at 37°C for 2 h with shaking in a humidified atmosphere. The pronuclei were then washed once in 1x DpnII buffer (New England Biolabs) with 1x BSA (New England Biolabs) and further digested overnight with 5 U DpnII (New England Biolabs) at 37°C in a humidified atmosphere. After a brief PBS wash and a wash through 1x ligation buffer (Thermo Scientific), the pronuclei were then ligated with 5 U T4 ligase (Thermo Scientific) at 16°C for 4.5 h with rotation (50 rpm), followed by 30 min ligation at RT. Next, Whole-genome amplification was performed using the illustra GenomiPhi V2 DNA amplification kit (GE Healthcare). In brief, the pronuclei were transferred to 0.2 ml PCR tubes in 3 μl sample buffer covered with mineral oil (Sigma-Aldrich) and were decrosslinked at 65°C overnight. Then, the pronuclei were lysed by adding 1.5 μl lysis solution (600 mM KOH; 10 mM EDTA; 100 mM DTT) and incubate for 10 min at 30°C, followed by neutralization with the addition of 1.5 μl neutralization solution (4 vol 1 M Tris HCl, pH 8.0; 1 vol 3M HCl). Whole genome amplification was carried out by addition of 4 μl samples buffer, 9 μl reaction buffer and 1 μl enzyme mixture and incubation at 30°C for 4 h followed by heat activation at 65°C for 10 min. High molecular weight DNA was purified using AMPure XP beads (Beckman Coulter, 1.8:1.0 beads:DNA ratio) and 1 μg DNA was sonicated to ~300-1300 bp fragments using the E220 Focused-ultrasonicator (Covaris). The sonicated DNA was purified with a PCR purification kit (Qiagen) and used to prepare Illumina libraries with the NEB Next Ultra Library Prep kit (Illumina). Libraries were sequenced on the HiSeq 2500 v4 with 125 bp paired-end reads (at VBCF NGS Unit) or on the NextSeq high-output lane with 75 bp paired-end reads (at MPIB NGS Core facility).

### snHi-C data analysis

snHi-C data were processed and analyzed similarly as in (28) and previously described in (2,31). Briefly, the reads of each sample were mapped to the mm9 genome with *bwa* and processed by the *pairtools* framework (https://pairtools.readthedocs.io/en/latest/) into pairs files. These data were subsequently converted into *COOL* files by the *cooler* package and used a container for Hi-C contact maps.

Loops were analyzed by summing up snHi-C contact frequencies for loop coordinates of over 12,000 loops identified using the Hi-C data from wildtype MEFs published in (31).We removed the effect of distance dependence by averaging 20 × 20 matrices surrounding the loops and dividing the final result by similarly averaged control matrices. Control matrices were obtained by averaging 20 × 20 matrices centered on the locations of randomly shifted positions of known loops (shifts ranged from 100 to 1,100 kb with 100 shifts for each loop). For display and visual consistency with the loop strength quantification, we set the backgrounds levels of interaction to 1. The background is defined as the top left 6 × 6 and the bottom right 6 × 6 submatrices. To quantify the loop strength, the average signal in the middle 6 × 6 submatrix is divided by the average signal in the top left and bottom right (at the same distance from the main diagonal) 6 × 6 submatrices.

For average TAD analysis, we used published TAD coordinates for the CH12-LX mouse cell line (45). We averaged Hi-C maps of all TADs and their neighbouring regions, chosen to be of the same length as the TAD, after rescaling each TAD to a 90 × 90 matrix. For visualization, the contact probability of these matrices was rescaled to follow a shallow power law with distance (-0.25 scaling). TAD strength was quantified using contact probability normalized snHi-C data. In Python notation, if M is the 90 × 90 TAD numpy array (where numpy is np) and L = 90 is the length of the matrix, then TAD_strength = box1/box2, where box1 = 0.5 * np.sum(M[0:L//3, L//3:2*L//3]) + 0.5 * np.sum(M[L//3:2*L//3,2*L//3:L]); and box2 = np.sum(M[L//3:2*L//3,L//3:2*L//3]).

To calculate the insulation score, we computed the sum of read counts within a sliding 40- kb-by-40-kb diamond. The diamond was positioned such that the “tip” touched the main axis of the snHi-C map corresponding to a “self-interaction”. Since snHi-C maps are not iteratively corrected, we normalized all insulation profiles by the score of the minimum insulation and then subtracted 1. This way, the insulation/domain boundary is at 0 and has a minimum of 0.

Contact probability P_c_(s) curves were computed from 10-kb binned snHi-C data. We divided the linear genomic separations into logarithmic bins with a factor of 1.3. Data within these logspaced bins (at distance, s) were averaged to produce the value of P_c_(s). Both P_c_(s) curves and their log-space slopes are shown following a Gaussian smoothing (using the scipy.ndimage.filters.gaussian_smoothing1d function with radius 0.8). Both the y-axis (i.e., log(P_c_(s)) and the x-axis (i.e., log[s]) were smoothed. The average loop size was determined by studying the derivative of the P_c_(s) curve in log-log space, that is, the slope of log(P_c_(s)). The location of the maximum of the derivative curve (i.e., position of the smallest slope) closely matches the average length of extruded loops.

### *In situ* bulk Hi-C

Hi-C was performed largely as described in (45) with minor modifications. Briefly, ~5×10^6^ HCT116 cells were crosslinked in 1% formaldehyde for 10 min at RT, snap-frozen and stored at −80°C. After permeabilization in lysis buffer (0.2% Igepal, 10 mM Tris-HCl pH 8.0, 10 mM NaCl, 1x Halt Protease inhibitor cocktail) nuclei were isolated in 0.3% SDS in NEBuffer 3 at 62°C for 10 min. SDS was quenched with 1% Triton X-100 at 37°C for 1 h, then the nuclei were pelleted and resuspended in 250 μl DpnII buffer with 600 U DpnII (New England Biolabs) at 37°C. After overnight digestion, 200 U DpnII were added followed by 2 h more incubation. Then, nuclei were spun down and resuspended in fill-in mix (biotin-14-dATP [Thermo Fisher Scientific], dCTP, dGTP and dTTP [Thermo Fisher Scientific], Klenow Polymerase [NEB], 1x NEB 2 buffer) for 1.5 h at 37°C with rotation. After ligation at RT for 4 h with T4 ligase (NEB), the nuclei were pelleted, resuspended in 200 μl H_2_O and digested with proteinase K for 30 min at 55°C in presence of 1% SDS. NaCl was added to a final concentration of 1.85 M before cross-links were reversed at 65°C overnight. After ethanol precipitation and a 70%-80% ethanol wash, DNA was resuspended in 10 mM Tris EDTA, transferred to a Covaris microTUBE (Covaris) and sheared to ~300-1300 bp fragments on the E220 Focused-ultrasonicator (Covaris). DNA was then bound to Dynabeads MyOne Streptavidin C1 beads (Thermo Fisher Scientific) for biotin pull down. Beads were resuspended in H_2_O used for library preparation with the NEBNext Ultra II Library Prep kit for Illumina (NEB). Beads were then washed 4 times using Tween wash buffer (5 mM Tris-HCl, 1 M NaCl, 0.5 mM EDTA, 0.05% Tween20) and DNA was eluted using 95 % formamide, 10 mM EDTA at 65 °C for 2 min. After precipitation, DNA was washed with 70-80% EtOH and resuspended in H_2_O. The finished libraries were sequenced on the NovaSeq 6000 system with 100 bp paired-end reads (at VBCF NGS Unit) or on the NextSeq high-output lane with 75 bp paired-end reads (at MPIB NGS Core facility).

### In situ bulk Hi-C data analysis

Bulk Hi-C data processing (both for data we generated and published data from ((46)) was performed using *distiller* - a nextflow based pipeline (https://github.com/open2c/distiller-nf, (47)). Reads were mapped to the hg38 reference genome with default settings except *dedup/max_mismatch_bp=0*. Multiresolution cooler files (48) generated by *distiller* were used for visualization in HiGlass (49) and in the downstream analyses.

For downstream analysis, we used *quaich* (https://github.com/open2c/quaich), a new snakemake pipeline for Hi-C postprocessing. It uses *cooltools*

(https://github.com/open2c/cooltools,(50)), *chromosight* (51) and *coolpup.py (52)* to perform compartment and insulation analysis, peak annotation and pileups, respectively. The config file we used is available here:https://gist.github.com/Phlya/5c2d0688610ebc5236d5aa7d0fd58adb.

We annotated peaks of enriched contact frequency in untreated HCT116 cell from (46) using *chromosight* (51) at 5 kbp resolution with default parameters. Then we used this annotation to quantify strength of Hi-C peaks in our datasets using pileups at 5 kbp resolution. Similarly, valleys of insulation score at 10 kbp resolution with 500 kbp window (and prominence over 0.1) were identified in the same published dataset and filtered to remove those that don’t disappear upon cohesin depletion (or don’t become at least 5-fold weaker) to identify cohesin-dependent domain boundaries. These were used to quantify changes in insulation in our datasets. Neighbouring insulation valleys were joined together to form TADs; regions longer than 1.5 Mbp were ignored. TAD coordinates were used for rescaled pileup analysis (28) to quantify their strength in our datasets.

### RNA-Seq G1 zygotes

For each replicate, a pool of 10 G1 zygotes were lysed, total RNA was extracted, and cDNA was synthesized using the SMART-Seq v4 Ultra Low Input RNA Kit (Takara Bio Europe). Sequencing libraries were prepared with the Nextera^®^ XT DNA Library Preparation Kit for Illumina^®^. Libraries were sequenced on the HiSeq 2500 v4 with 50 bp single-end reads at VBCF NGS Unit.

### RNA-Seq tissue culture cells

Total RNA from HCT116 cells were isolated using a lysis step based on guanidine thiocyanate (adapted from (53) and using magnetic beads (GE Healthcare, 65152105050450)). mRNA sequencing libraries were prepared from 1 μg total RNA using NEBNext^®^ Poly(A) mRNA Magnetic Isolation Module (E7490) and NEBNext^®^ Ultra™ II Directional RNA Library Prep Kit for Illumina^®^ (E7760). Paired-end sequencing was performed on Illumina NextSeq 500 (2 × 43 bp reads). Total 9 samples were multiplexed and sequenced on a NextSeq 500/550 High Output Kit v2.5 (75 Cycles) at MPIB NGS Core facility. BCL raw data were converted to FASTQ data and demultiplexed by bcl2fastq.

### RNA-Seq analysis

FASTQ files from sequencing mouse G1 zygotes or human HCT116 cell line were pseudoaligned to mm10 or hg38 releases of Mus musculus and Homo sapiens genomes, respectively using Kallisto with 100 bootstraps (54). Resulting abundance measures were analysed in R to generate PCA plots (55) and heatmap of the correlation matrix (56). To find differentially expressed transcripts we used Wald test for Sleuth model (sleuth) in R (57).

Changes in chromatin contact frequencies were analysed by aggregating the number of contacts, normalised with HiCcompare in R (58), around DE and non-DE TSS observed in DMSO or auxin treated MCM-AID expressing HCT116 cells in distance bins of 0-10 kbp, 10-100 kbp, 100-500 kbp and over 500 kbp using HiC maps of 10 kbp resolution. Mean change of contact frequencies in each bin for every category was calculated by averaging the auxin vs DMSO treatment ratios of normalised sum of contacts. All the mean contact frequency changes were tested against the non-DE TSS control using the non-parametric Kruskal-Wallis test followed by pairwise Wilcox (Mann Whitney U) test.

All plots were compiled with ggplot2 in R.

### Protein expression and purification

#### Cohesin

Human recombinant cohesin^STAG1, SCC1^-^Halo^ was purified and fluorescently labeled with Janelia Fluor 549 HaloTag (Promega) as previously described (5).

#### ORC and Cdc6

*S. cerevisiae* recombinant ORC and Cdc6 were purified as previously described (59).

#### SFP synthase

SFP synthase was purified essentially as previously described (60).

#### Cdt1-Mcm2-7

To generate fluorescently labeled *S. cerevisiae* recombinant Cdt1-Mcm2-7, *S. cerevisiae* strain ySA4 was generated. In brief, a ybbR and 3xFLAG tag were fused to the N- and C-terminus of Mcm6, respectively, generating Cdt1-Mcm2-7^ybbR-Mcm6^. Cells were grown in 6 L YP supplemented with 2 % (v/v) raffinose at 30 °C. At OD600 = 1.2, cells were arrested at G1 by adding α-factor to a final concentration of 150 ng/ml for 3 hours. Subsequently, protein expression was induced by addition of 2 % (v/v) galactose. After 4 hours, cells were harvested and washed once with cold MilliQ water + 0.3 mM PMSF and once with buffer A (100 mM HEPES-KOH, pH 7.6, 0.8 M Sorbitol, 10 mM Mg(OAc)2, 0.75 M potassium glutamate (KGlu)). Finally, cells were resuspended in 1 packed cell volume of buffer A + 1 mM DTT supplemented with a protease inhibitor cocktail (2 μM pepstatin, 2 μM leupeptin, 1 mM PMSF, 1 mM benzamidine, 1 μg/ml aprotinin) and frozen dropwise in liquid N2. Frozen cells were lysed in a freezer mill (SPEX) and lysed cell powder was resuspended in 1 packed cell volume buffer B (45 mM HEPES-KOH, pH 7.6, 0.02 % (v/v) Nonidet P40 Substitute, 5 mM Mg(OAc)2, 10 % (v/v) glycerol, 1 mM ATP, 1 mM DTT) + 300 mM KGlu. All subsequent purification steps were performed at 4 °C unless stated otherwise. The lysate was cleared by ultracentrifugation at 235000 g for 60 min. Soluble lysate was incubated with 0.5 ml bed volume (BV) Anti-FLAG M2 affinity gel (Sigma) equilibrated with buffer B + 300 mM KGlu for 3 h. The resin was washed twice with 20 BV buffer B + 300 mM KGlu and twice with 20 BV buffer B + 100 mM KGlu. Protein was eluted with buffer B + 100 mM KGlu + 0.5 mg/ml 3xFLAG peptide.

For site specific labeling, Cdt1-Mcm2-7^ybbR-Mcm6^ was incubated with SFP-Synthase and LD655-CoA (Lumidyne Technologies) at a 1:3:6 molar ratio for 2 h at 30 °C in buffer B + 100 mM KGlu, 10 mM MgCl2. Labeled protein was further purified on a Superdex 200 increase 10/300 gel filtration column (GE Healthcare) equilibrated in buffer B + 100 mM potassium acetate (KOAc). Protein containing fractions were pooled, concentrated with a MWCO 50000 Amicon Ultra Centrifugal Filter unit (Merck) and stored in aliquots at −80 °C. The labeling efficiency was estimated to be ~90 % from the extinction coefficients of Cdt1-Mcm2-7 and LD655.

#### Single-molecule imaging

Single-molecule assays were performed on a custom-built micromirror TIRF microscope similar as previously described (61) with an Apo N TIRF 60 × oil-immersion TIRF objective (NA 1.49, Olympus). Janelia Fluor 532 and LD655 were excited with a 532 nm and 637 nm laser (OBIS 532nm LS 120mW and OBIS 637 nm LX 100 mW, Coherent), respectively at a frame rate of ~6 fps. Residual scattered light from excitation was removed with a ZET532/640m emission filter (Chroma). Emission light was split at 635 nm (T635lpxr, Chroma) and recorded as dual-view with an iXon Ultra 888 EMCCD camera (Andor). All microscope parts were controlled using Micromanager v1.4 (62) and custom Beanshell scripts.

#### Preparation of PEG-Biotin microscope slides

Glass coverslips (22 × 22 mm, Marienfeld) were cleaned in a plasma cleaner (Zepto, Diener Electronic) and subsequently incubated in 2 % (v/v) 3-aminopropyltri ethoxy silane (Roth) in acetone for 5 min. Silanized coverslips were washed with ddH_2_O, dried and incubated at 110 °C for 30 min. Slides were covered with a fresh solution of 0.1 M NaHCO_3_ containing 0.4 % (w/v) Biotin-PEG-SC-5000 and 15 % (w/v) mPEG-SC-5000 (Laysan Bio) and incubated overnight. Functionalized slides were washed with ddH2O, dried and incubated again overnight in a fresh Biotin-PEG/mPEG solution. Slides were finally washed, dried and stored under vacuum.

#### DNA substrate for single-molecule imaging

To generate pMSuperCos-ARS1, first, a 21 kb genomic DNA fragment of bacteriophage lambda (NEB) was flanked by a unique XbaI (position 0) and NotI restriction site on either end and cloned into pSuperCos1 backbone (Stratagene). Second, yeast origin ARS1 was inserted at BamHI site around position 5.3 kb within the 21 kb genomic DNA fragment.

To produce the DNA substrate for single-molecule imaging, pMSuperCos-ARS1 was isolated from DH5α using a Plasmid Maxi Kit (Qiagen). 100 ug plasmid were digested with 100 U Notl-HF and XbaI (NEB) for 7 h at 37 °C. The resulting 21202 bp ARS1-DNA fragment was separated from the SuperCos1 backbone on a 10-40% sucrose gradient. DNA handles were prepared by annealing oligonucleotides MS_200/201 MS202/203 in equimolar amounts in 30 mM HEPES, pH 7.5, 100 mM KOAc by heating to 95 °C for 5 min and cooling to 4 °C at −1 °C/min. Annealed handles were mixed with the purified 21 kb ARS1-DNA at a molar ratio of 15:1 and ligated with T4 DNA Ligase in 1 × T4 ligase buffer (both NEB) at 16 °C overnight. Free handles were removed on a Sephacryl S-1000 SF Tricorn 10/300 gel filtration column (GE Healthcare) equilibrated in 10 mM Tris, pH 8, 300 mM NaCl, 1 mM EDTA. Peak fractions were pooled, EtOH precipitated and reconstituted in TE buffer. Final DNA was stored in aliquots at −80 °C. Note that the final linear DNA is functionalized with biotin at NotI site and an 18 bp ssDNA overhang at XbaI site which is used for orientation specific doubly-tethering.

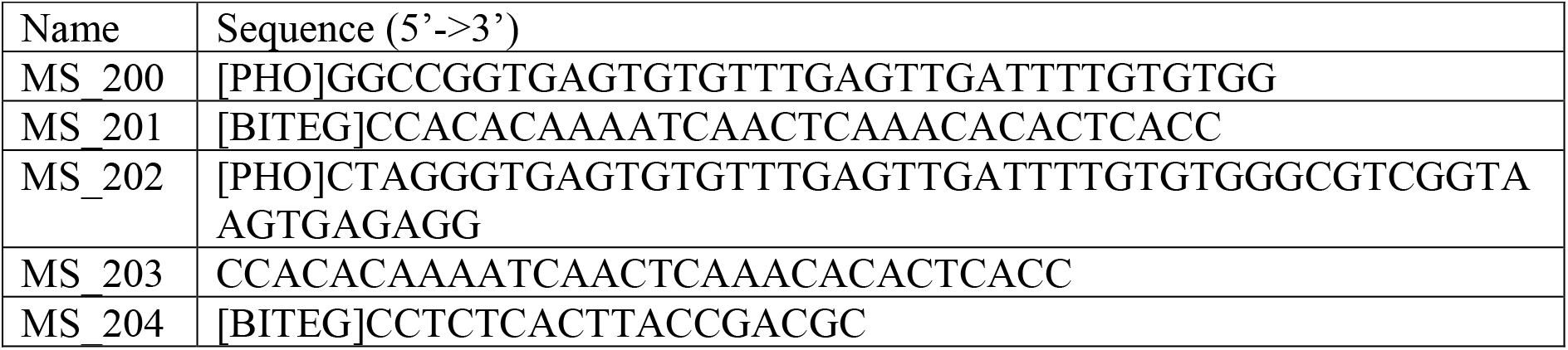
Oligonucleotide sequences. All oligonucleotides were synthesized by Eurofins Genomics (Ebersberg, Germany).

#### Flow cell preparation

A functionalized PEG-Biotin slide was incubated with blocking buffer (20 mM Tris-HCl, pH 7.5, 50 mM NaCl, 2 mM EDTA, 0.2 mg/ml BSA, 0.025 % (v/v) Tween20) + 0.2 mg/ml streptavidin (Sigma) for 30 min. A flow cell was assembled by placing a polydimethylsiloxane block on top to generate a 0.5 mm wide and 0.1 mm high flow channel and a polyethylene tube (inner diameter 0.58 mm) was inserted on either end.

DNA was introduced to the flow cell at 5 pM in blocking buffer and incubated for 15 min in absence of buffer flow to allow binding to the slide surface. To doubly tether DNA, the flow lane was flushed with 100 uM oligonucleotide MS_204 in blocking buffer + 0.2 mM chloroquine (Sigma) at 100 ul/min.

#### Single-molecule sliding assay

Helicase loading was achieved by introducing 0.25 nM ORC, 4 nM Cdc6 and 10 nM Cdt1-Mcm2-7^ybbR-LD655-Mcm6^ in licensing buffer (30 mM HEPES-KOH, pH 7.6, 8 mM Mg(OAc)2, 0.1 mg/ml BSA, 0.05 % (v/v) Tween20) + 200 mM KOAc, 5 mM DTT, 3 mM ATP to a prepared flow cell and incubating for 25 min. Cohesin loading and sliding was essentially performed as previously described (38). 0.7 nM cohesin^STAG1,SCC1-Halo-JF546^ were incubated with licensed DNA in cohesin binding buffer (35 mM Tris, pH 7.5, 25 mM NaCl, 25 mM KCl, 1 mM MgCl2, 10 % (v/v) glycerol, 0.1 mg/ml BSA, 0.003 (v/v) Tween20, 1 mM DTT, 0.2 mM ATP) for 10 min. Unbound protein was washed out and cohesin sliding was induced by changing to licensing buffer + 500 mM NaCl, 1 mM DTT, 0.6 mM ATP supplemented with an oxygen scavenging system (1 mM Trolox, 2.5 mM PCA, 0.21 U/ml PCD (all Sigma)) (63) and imaging was started. DNA was post stained with 50 nM Sytox Orange (Thermo Fisher) in the same buffer as during imaging.

#### Single-molecule data analysis

Single-molecule data were analyzed using Fiji, Molecule ARchive Suite (MARS) Fiji plugin (https://github.com/duderstadt-lab/) and custom python scripts. Briefly, all doubly-tethered DNA molecules containing cohesin were chosen for analysis. Cohesin and MCM were tracked individually and merged with DNA to determine their position on the same DNA molecule.

The probability of cohesin passing MCM was addressed as follows: Frames in which cohesin colocalized with MCM (mean position) within less than 1.5 kb were classified as encounter. Upon an encounter, if cohesin passed MCM in the consecutive frame by at least 0.5 kb, the encounter was determined as successful passing. All remaining frames (distance > 1.5 kb to MCM) were further evaluated for MCM passing as described above and additionally counted as encounter with successful passing. DNA molecules with cohesin only were analyzed the same way using the theoretical ARS1 position on DNA.

MCM photobleaching steps were defined as abrupt drops in fluorescence intensity and detected using the kinetic change point analysis (64).

Diffusion coefficients (D) were calculated with:

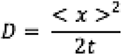

with < x >^2^ being the mean square displacement in kbp^2^ and t the time in s.

All kymographs were generated using Fiji. For this, individual DNA ends were fitted with subpixel localization and the kymograph was generated along the connecting line. Individual DNA molecules doubly-tethered with different extension to the slide surface and as a consequence, kymographs differ in heights. These length differences were accounted for throughout all analysis steps described above.

### Loop extrusion simulations and contact map generation

#### Time steps and lattice set-up

We use a fixed-time-step Monte Carlo algorithm as in previous work (39). We define the chromosome as a lattice of *L*= 10,000 sites, where each lattice site corresponds to 2 kb of DNA. Loop extruding factors (LEFs) are represented as two motor subunits, which move bidirectionally away from one another one lattice site at a time. When LEFs encounter one another, we assume that they cannot bypass each other as is typical for cohesin simulations (65). The ends of the chromosome (i.e. the first and last lattice sites) are considered boundaries to LEF translocation; this way, LEFs cannot “walk off” the chromosome.

#### CTCF and MCM boundary elements

To simulate TADs, we specify that every 150^th^ lattice site is a CTCF site. In this way, our simulated 20 Mb chromosome segment is comprised of 66 TADs each of size 300 kb. CTCF sites may stall translocation of a LEF subunit with a probability of 0.45. This stalling probability is chosen within the experimental estimates of 15%-50% fractional occupancy of CTCF sites via ChIP-seq and microscopy (66). For simulations mimicking the “control” and “Wapl” depletion conditions (i.e. where MCM is present on the genome), we also add random extrusion barriers to our lattice to mimic the presence of MCMs. For our parameter sweep, we add 33, 66, 132, 264, 528 barriers (i.e. representing MCMs) randomly dispersed in the 20 Mb chromosome segment; this corresponds to a density of 1 MCM complex per 600 kb, 300 kb, 150 kb, 75 kb, 37.5 kb, respectively. The MCM barriers are fixed in place for the duration of a simulation. Like the CTCFs, the MCM barriers can also stall LEF translocation. A randomly translocating LEF subunit will be stalled at an MCM site with a probability of 0.0001, 0.05, 0.2, 0.4 or 0.8 (i.e. meaning that LEFs can bypass between ~20-100% of MCM sites). For both CTCF and MCM lattice sites, “stalling” a LEF subunit is a permanent event which prevents further movement of that subunit. Stalling events are only resolved upon dissociation of the LEF from the lattice. For simulations where there is ‘MCM loss’, we set the total number of random MCM barriers to zero but keep the CTCF lattice sites the same. All results presented in this paper are from an average over 25 different random distributions of MCMs (i.e. 25 simulation runs were performed for each condition).

#### LEFs separations and processivity

For our simulations of ‘control’ and ‘MCM loss’ conditions, the default LEF processivity was 90 kb, and the default LEF separation was 300 kb. For our simulations of the *‘Wapl^∆^’* and ‘*Wapl^∆^*/MCM loss’ conditions, the LEF processivity was 130 kb, and the separations were 180 kb. The ~50% increase in density after Wapl depletion is supported by quantitative immunofluorescence data indicating there is a modest enrichment of cohesin after removal of Wapl (35).

#### Association and dissociation rates

All simulations are performed with fixed numbers of extruders, which results in equal association and dissociation rates. The dissociation rate is ultimately tied to the “processivity” of the LEF, which is the average distance in kb (or lattice sites) which the LEF travels before dissociating. We allow LEFs to randomly associate to at any lattice position after a dissociation event.

#### Loop extrusion equilibration steps

We compute 10,000 initialization steps for each simulation before creating any contact maps. This ensures that the loop statistics have reached a steady-state. Subsequent loop configurations were sampled every 100 simulation steps in order to generate contact maps. We sampled from at least 2,500 different LEF configurations (i.e. 100 configurations from 25 different simulations) to generate contact probability decay curves and perform aggregate peak analysis (see below).

#### Contact maps

We generated contact maps semi-analytically, which employs a Gaussian approximation to calculate contact probability maps directly from the positions of LEFs. This approach was developed previously (39) and used to simulate bacterial Hi-C maps. We note that since the density of cohesins is sufficiently low in the zygotes (i.e. the processivity and separation ratio is close to or less than 1), and since the contact probability scaling exponent up to 10 Mb is close to −1.5 in the absence of cohesins (2), we are justified in using the Gaussian approximation to generate contact maps. To generate the P_c_(s) curves, we utilize at least 9,000,000 random samples of the contact probability; these samples were taken from varying genomic positions and relative separations within the simulated 20 Mb of chromosome and averaged using logarithmically spaced bins (factor of 1.3). To generate the equivalent of the aggregate peak analysis for contact enrichments at CTCF sites, we utilized at least 144,000,000 random samples of the contact probability from a 100 kb by 100 kb window centered on the CTCF sites. These 144,000,000 samples were distributed evenly between 64 TADs (there are 66 TADs, but we excluded the two TADs closest to the chromosome ends) and at least 2,500 LEF conformations. Control matrices for normalization were obtained as described above, but using a shifted window shifted by 150 kb from the TAD boundaries. Aggregate peak analysis plots are shown coarse-grained to 20×20 bins.

#### Estimation of chromatin-bound MCM density in mammalian cells

Using mass-spectrometry analysis, the copy number of each MCM subunit is estimated at ~670.000 in HeLa cells (67), and quantitatively immunoblotting shows that in late G1 phase ~45% of MCM2 is bound to chromatin (68). This leads to the estimate that ~301.500 MCMs are bound to the chromatin in late G1. Knowing that MCMs form double hexamers on chromatin and that the average genome size of HeLa cells is ~7.9 × 10^9^ (69), we estimate a density of 1 MCM double hexamer every ~52 kb (7.9 × 10^9^/(301500/2)) (assuming a random distribution of MCMs).

